# Modulation of tooth regeneration through opposing responses to Wnt and BMP signals in teleosts

**DOI:** 10.1101/2022.10.01.510447

**Authors:** Tyler A. Square, Emma J. Mackey, Shivani Sundaram, Naama C. Weksler, Zoe Z. Chen, Sujanya Narayanan, Craig T. Miller

## Abstract

Most vertebrate species undergo tooth replacement throughout adult life. This process is marked by the shedding of existing teeth and the regeneration of tooth organs. However, little is known about the genetic circuitry regulating tooth replacement. Here we tested whether fish orthologs of genes known to regulate mammalian hair regeneration have effects on tooth replacement. Using two fish species that demonstrate distinct modes of tooth regeneration, threespine stickleback (*Gasterosteus aculeatus*) and zebrafish (*Danio rerio*), we found that transgenic overexpression of four different genes changed tooth replacement rates in direction predicted by a hair regeneration model: *Wnt10a* and *Grem2a* increased tooth replacement rate, while *Bmp6* and *Dkk2* strongly inhibited tooth formation. Thus, similar to known roles in hair regeneration, Wnt and BMP signals promote and inhibit regeneration, respectively. Regulation of total tooth number was separable from regulation of replacement rates. RNA-seq on stickleback dental tissue showed that *Bmp6* overexpression resulted in an upregulation of Wnt inhibitors. Together these data support a model where different epithelial organs like teeth and hair share genetic circuitry driving organ regeneration.

## Introduction

Organs are often replaced or renewed throughout the life of an organism as a feature of wild-type development. These programmed regenerative processes are especially common in epithelia and their associated organs, like teeth (Tucker and Fraser, 2014; Jernvall and Thesleff, 2012; Richman et al., 2013). Tooth replacement usually takes the form of whole tooth regeneration, where entire new tooth organs differentiate while old teeth are removed by active shedding and/or dislodgement through use. Within species, tooth replacement events often form characteristic patterns, exhibiting a high degree of consistency with respect to their timing, sequence, and spacing (Berkovitz and Shellis, 2023; Whitlock and Richman, 2013). This consistency has led to hypotheses that neighboring or alternating tooth positions could be subject to signals that act to influence or coordinate the timing of replacement cycles for multiple tooth positions in tandem (DeMar, 1972; Edmund, 1960; Fraser and Thiery, 2020; Grieco and Richman, 2018; Osborn, 1970; Osborn, 1998). The potential involvement of secreted signals in tooth replacement is additionally suggested by studies on primary tooth differentiation, where the functional relevance of numerous signaling ligands has been documented (Fraser et al., 2013; Tucker and Sharpe, 2004; Yu and Klein, 2020). However, little is known about which specific ligands can regulate the process of whole tooth replacement.

Whole organ replacement is a trait that teeth share with other epithelial appendages. This class of organs includes body coverings like hair, scales, and feathers, and some soft organs like salivary and sweat glands (Chuong and Noveen, 1999; Chuong et al., 2012; Wu et al., 2004). Whether regeneration is cyclic and programmed, or brought on by injury or wear, nearly all types of epithelial appendages undergo whole-organ regeneration (Cox et al., 2018; Finger and Barlow, 2021; Lin et al., 2013; Lin et al., 2021; Plikus et al., 2008; Rocchi et al., 2021). Despite the stark differences in their basic compositions as mature organs, different epithelial appendages demonstrate numerous developmental genetic similarities, in some cases suggesting deep homology or even direct homology between these organs (Aman et al., 2018; Benton et al., 2019; Debiais-Thibaud et al., 2011; Di-Poï and Milinkovitch, 2016; Martin et al., 2016; Pispa and Thesleff, 2003; Rasch et al., 2016; Sharpe, 2001; Wu et al., 2004). Given that most epithelial appendages exhibit whole-organ regeneration, we parsimoniously hypothesize that these regenerative processes are driven by shared genetic networks. In support of this hypothesis, we previously documented the expression of ten candidate hair follicle stem cell marker gene orthologs in both zebrafish and stickleback successional dental epithelia (SDE), including *Bmp6*, *CD34, Nfatc1*, *Lgr6*, and *Gli1* (Square et al., 2021), suggesting a remarkable level of genetic overlap in these naïve epithelial tissues.

Here we aimed to test whether genes implicated in hair regeneration could similarly influence tooth regeneration in two teleost models: threespine stickleback (*Gasterosteus aculeatus*) and zebrafish (*Danio rerio*). In mammalian hair, some secreted factors have been identified as likely or possible regulators of hair follicle stem cells. Namely, a Wnt-BMP cycling mechanism has been well supported, whereby the Wnt and BMP pathways have oppositional roles that drive the oscillation of the hair regenerative cycle between active growth (anagen, high Wnt + BMP inhibitors) and quiescence (telogen, high BMP + Wnt inhibitors) (Daszczuk et al., 2020; Kandyba et al., 2013; Plikus et al., 2008; Wu et al., 2019). Ligands implicated in promoting hair regeneration include the Wnt ligands Wnt10a, Wnt10b, Wnt7a, and Wnt7b, as well as the BMP inhibitors Grem1, Grem2, Bambi, and Noggin1; conversely, secreted BMP signals like Bmp2, Bmp4, Bmp6, and Wnt inhibitors like Dkk1 and Dkk2 have been implicated in slowing or stopping the regenerative process (Xu et al., 2017; Kandyba et al., 2013; Wu et al., 2019; Niiyama et al., 2018; Niiyama et al., 2022; Botchkarev et al., 2001; Greco et al., 2009; Kishimoto et al., 2000; He et al., 2022; Harshuk-Shabso et al., 2020; Kwack et al., 2012; Plikus et al., 2008; Rendl et al., 2008). To test whether such secreted ligands could elicit congruous changes in the replacement rates of teeth, we selected coding sequences for four of the above ligands (*Wnt10a*, *Dkk2*, *Bmp6*, and *Grem2a*) to test for both endogenous expression in actively regenerating tooth fields, and their possible effects on tooth replacement rates and/or total tooth number. Our selection of gene orthologs was motivated by known pleiotropic disease loci in humans and genetic studies from other vertebrates. *WNT10A* and *GREM2* are known to be associated with different forms of ectodermal dysplasia, wherein both tooth and hair regeneration are perturbed, but not always primary epithelial organ growth (Mostowska et al., 2018; Xu et al., 2017). *Grem2* loss-of-function has additionally been shown to disrupt constant incisor outgrowth in mice (Vogel et al., 2015). *Bmp6* has been strongly implicated in both mouse hair regeneration (Kandyba et al., 2013; Wu et al., 2019) and the natural evolution of stickleback tooth replacement rates and total tooth number (Cleves et al., 2014; Cleves et al., 2018). *Dkk2* expression has been shown to oscillate during the regenerative cycle in the hair follicle (Harshuk-Shabso et al., 2020) and disruption of this gene in mice causes ectopic hair to grow in normally hairless regions of skin (Song et al., 2018). Transcripts from all four of these selected genes were detected in previously published RNA-seq datasets derived from late-stage stickleback tooth fields undergoing replacement (Hart et al., 2018; Mack et al., 2023), suggesting these genes could regulate tooth replacement. Furthermore, stimulation of downstream Wnt signaling via a constitutively active β-catenin (*Ctnnb*) has been shown to induce regeneration events in mouse molars, a process which normally does not occur in this species, supporting a crucial role for Wnt signaling in promoting whole tooth organ regeneration (Järvinen et al., 2006; Popa et al., 2019).

## Results

### Expression of secreted ligand genes of interest in sticklebacks

We examined *Wnt10a*, *Dkk2*, *Bmp6*, and *Grem2a* expression in wild-type subadult stickleback pharyngeal tooth fields (Fig. 1). We documented expression not just in tooth organs themselves, but also in the regions between teeth, as these inter-tooth expression domains could also be involved with regulating tooth organ development or regeneration. Overall, we found that all four genes are expressed both in developing tooth organs and in epithelial and/or mesenchymal cell populations surrounding tooth organs.

**Figure 1.**
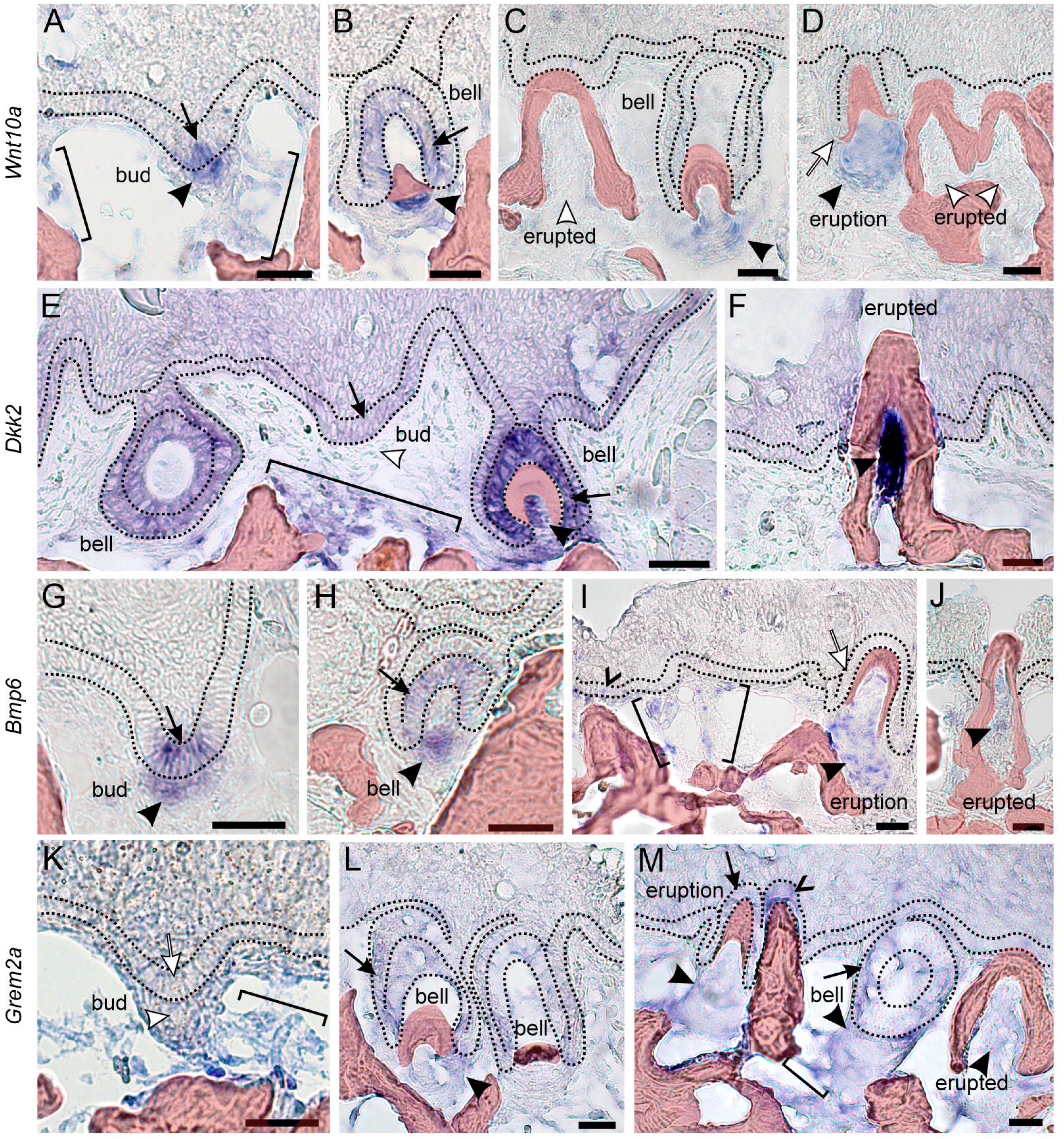
*In situ* hybridization reveals dynamic expression of *Wnt10a*, *Dkk2*, *Bmp6*, and *Grem2a* within and surrounding pharyngeal teeth in wild-type sticklebacks. The basalmost layer of epithelium is flanked by black dotted lines in each image. Arrows mark dental epithelium, arrowheads mark dental mesenchyme, black pointers mark detected expression, white markers indicate regions with no detected expression, brackets mark mesenchymal expression outside of tooth organs, and carets mark the successional dental epithelium. Bony tissues are false-colored red. Tooth stages and genes are labeled in the figure. Each expression pattern presented was observed in at least n=3 individuals. Scale bars: 20 μm.

*Wnt10a* transcripts were detected in early bud-stage tooth germs in both the epithelium and the earliest tooth mesenchyme (Fig. 1A, black arrow and arrowhead). Epithelial expression was also observed at the mid-bell stage, favoring the inner dental epithelium (Fig. 1B, black arrow), but was not appreciably detected in later bell stage tooth germ epithelium (Fig. 1C,D, white arrows). *Wnt10a* expression was also found in the dental mesenchyme of bell and eruption stage teeth (Fig. 1B-D, black arrowheads) but was not appreciably detected in fully ankylosed and erupted teeth (Fig. 1C,D, white arrowheads). *Wnt10a* was also detected in isolated regions of epithelial and mesenchymal cells at other locations in the tooth field near or below developing tooth germs (Fig. 1A, brackets).

The Wnt signaling inhibitor *Dkk2* was diffusely expressed in the epithelium overlying the entire tooth field (Fig. 1E, F). Marked epithelial expression was detected in bud and bell-stage tooth organs, especially in the inner dental epithelium of bell stages (Fig. 1E, black arrows). In tooth mesenchyme, we did not detect expression in bud stage tooth germs (Fig. 1E, white arrowhead), but strong expression was observed at the early bell stage (Fig. 1E, black arrowhead) and across more advanced tooth stages, especially in odontoblasts near the tooth apex (tip) in fully erupted and ankylosed teeth (Fig. 1F, black arrowhead). *Dkk2* was additionally observed in deep mesenchymal cell populations between teeth, against the bone of attachment that serves as the anchor for ankylosed teeth (Fig. 1E, bracket).

*Bmp6* was expressed similarly to *Wnt10a* in bud-stage teeth, exhibiting focal expression in both the epithelium and mesenchyme (Fig. 1G, black arrow and arrowhead). *Bmp6* was additionally detected in the inner dental epithelium and mesenchyme of bell-stage tooth germs (Fig. 1H, black arrow and arrowhead), and mesenchymal expression was observed in eruption stages and in fully erupted teeth (Fig. 1I, J, black arrowheads). *Bmp6* expression was also detected at a subset of successional dental epithelia (SDE, Fig. 1I, caret) as well as isolated clusters of mesenchymal cells surrounding teeth in the tooth field (Fig. 1I, brackets), though we did not detect it in eruption stage tooth epithelium (Fig. 1I, white arrow).

The BMP inhibitor *Grem2a* was not detected in bud stage teeth themselves (Fig. 1K, white arrow and arrowhead), though it was detected in cells surrounding the core of the bud stage tooth mesenchymal condensation (Fig. 1K, bracket). During bell stages, tooth germs exhibited both inner and outer dental epithelial expression as well as mesenchymal expression (Fig. 1L, M, black arrows and arrowhead), which appeared to persist in eruption stage teeth (Fig. 1M, black arrow and arrowhead). Fully ankylosed and erupted teeth also demonstrated mesenchymal expression in odontoblasts (Fig. 1M, black arrowheads). *Grem2a* transcripts were additionally detected in the SDE (Fig. 1M, caret) and diffusely in other dispersed epithelial cells (Fig. 1K-M).

### Pulse-chase bone labeling and gene overexpression approach

Three of the four secreted ligand genes we focus on here are known to be required for normal primary tooth development in various species: *Wnt10a* (Xu et al., 2017; Yuan et al., 2017)*, Bmp6* (Cleves et al., 2018), and *Grem2a* (Mostowska et al., 2018; Vogel et al., 2015). These early requirements for dental development present obstacles to using germline loss-of-function mutations to understand gene functions in late-stage tooth fields, because alterations to early dental differentiation likely alter later events like regeneration. We thus sought a temporally inducible genetic system that could test the effects of secreted Wnt and BMP ligands and inhibitors during late developmental stages without interfering with tooth field initiation and early primary tooth differentiation. Heat shock gene overexpression (OE) was therefore used here to induce transgenes in sub-adult and adult fish, a well-established method in zebrafish (Duszynski et al., 2011; Shoji and Sato-Maeda, 2008). We thus coupled OE treatments of *Wnt10a*, *Dkk2*, *Bmp6*, or *Grem2a* with a two-color pulse-chase bone staining protocol (Figs 2 and S1), allowing us to classify every tooth in each fish as either “new” or “retained” with respect to the OE treatment interval (see Methods). By calculating the new:retained tooth ratio for each fish, we describe a simple proxy of tooth turnover rates in each individual. We additionally summed all new and retained teeth to assess “total tooth number,” a count that includes both erupted functional teeth and unerupted bony tooth germs. In sticklebacks this count includes both oral and pharyngeal teeth usually numbering ∼200-300, whereas zebrafish have just one pair of pharyngeal tooth fields with ∼25-30 total teeth. Since zebrafish exhibit morphologically stationary tooth families with a stereotypical number (11) and arrangement that is reached during early juvenile stages (Van der Heyden and Huysseune, 2000), we additionally ask whether OE treatments can modify the number of tooth families present in zebrafish.

**Figure 2.**
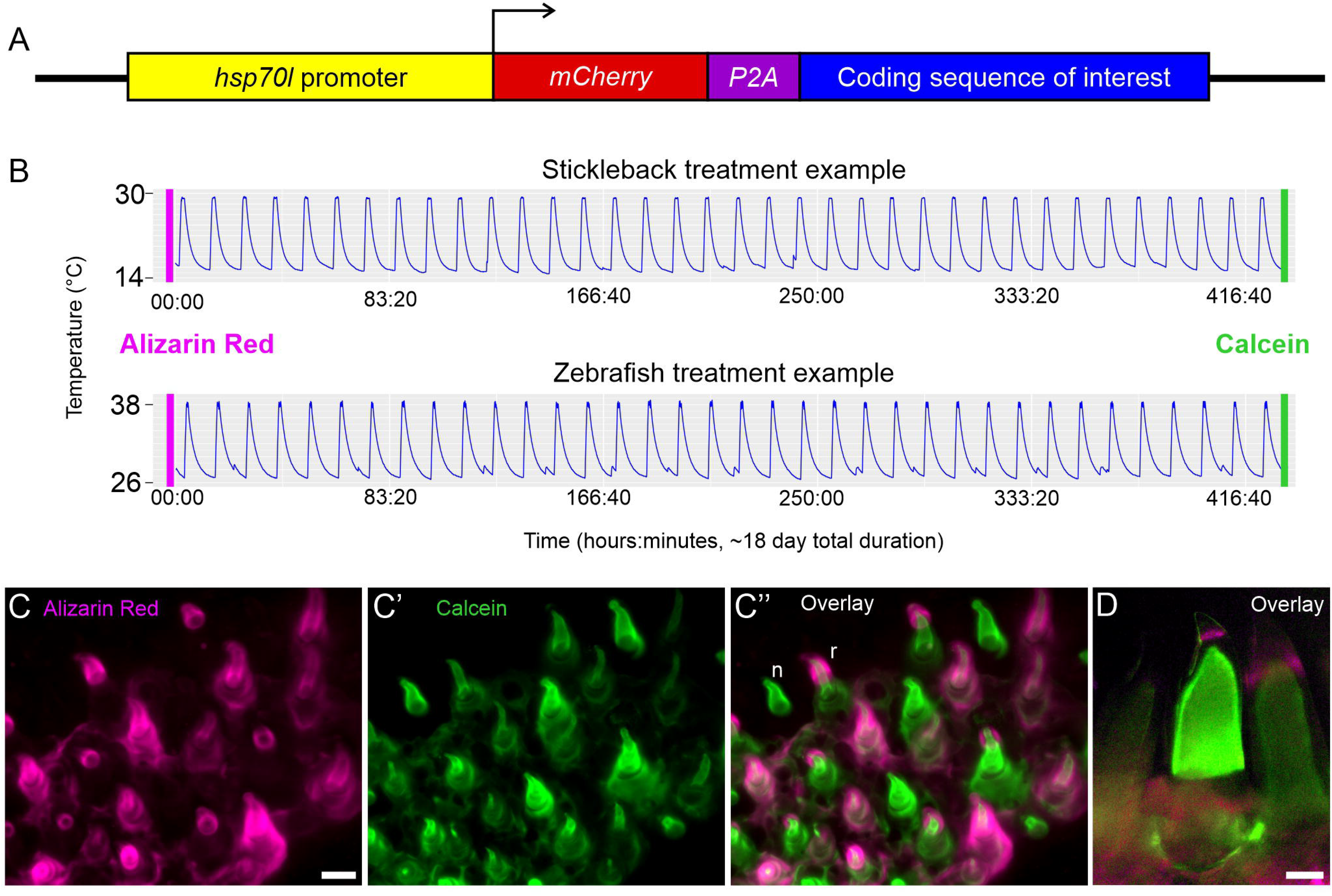
Summary of the two-stain pulse-chase heat shock method for assessing tooth gain and loss. **A.** Schematic of OE transgenes. The zebrafish *hsp70l* promoter drives an mCherry-P2A-CDS transgene, where “CDS” is one of seven full-length coding sequences of interest (see Methods and Table S2). P2A is a viral 22 amino acid sequence that cleaves between residues 20 and 21, separating mCherry from the CDS. **B.** Example temperature profiles for heat shock treatments in sticklebacks (top) and zebrafish (bottom). Alizarin pulse and calcein chase are shown as pink and green bars, respectively. **C.** Example images of the pulse-chase method on a control stickleback. Alizarin Red (F) strongly marks all bone undergoing active ossification at the start of the treatment (magenta). **C’.** 18 days later, after 36 heat shocks, a calcein chase marks all bone ossifying at the end of the treatment (green). **C’’.** Overlain images showing Alizarin Red and calcein revealed whether individual teeth were either present at only the 2^nd^ labeling (new, calcein only, example labeled “n”) or present at both the first and second labeling (retained, Alizarin Red and calcein positive, example labeled “r”). **D.** An overlay of the pulse-chase treatment on zebrafish teeth using the same treatment interval. Scale bars: 25 μm.

### Wnt pathway modulation by overexpression of *Wnt10a* and *Dkk2*

We first tested whether activation of the Wnt pathway via *Wnt10a* overexpression was associated with an increase in tooth replacement rates or total tooth number (Fig. 3). *Wnt10a* OE in sticklebacks (Fig. 3A) simultaneously increased the number of new teeth and reduced the number of retained teeth. Together, these two shifts consistently raised the average new:retained tooth ratio. However, *Wnt10a* OE did not significantly change the total number of teeth, indicating that increasing the replacement rate does not necessarily alter total tooth number. These shifts are also reflected by most tooth field types alone (Fig. S2). Qualitatively, we noticed that the *Wnt10a* OE individuals oftentimes displayed uninterrupted clusters of five or more new teeth (Fig. 3B,C, dotted oval), whereas new tooth distribution in wild-type (WT) controls appeared more uniform. Measuring the area of the ventral tooth plates (VTPs) showed that tooth field area was not significantly altered under *Wnt10a* OE (Fig. S3). We additionally carried out a negative control assay with this transgene to test for latent effects on tooth replacement in non-heat shocked transgene carriers; we found no such effects (see methods, Fig. S4).

**Figure 3.**
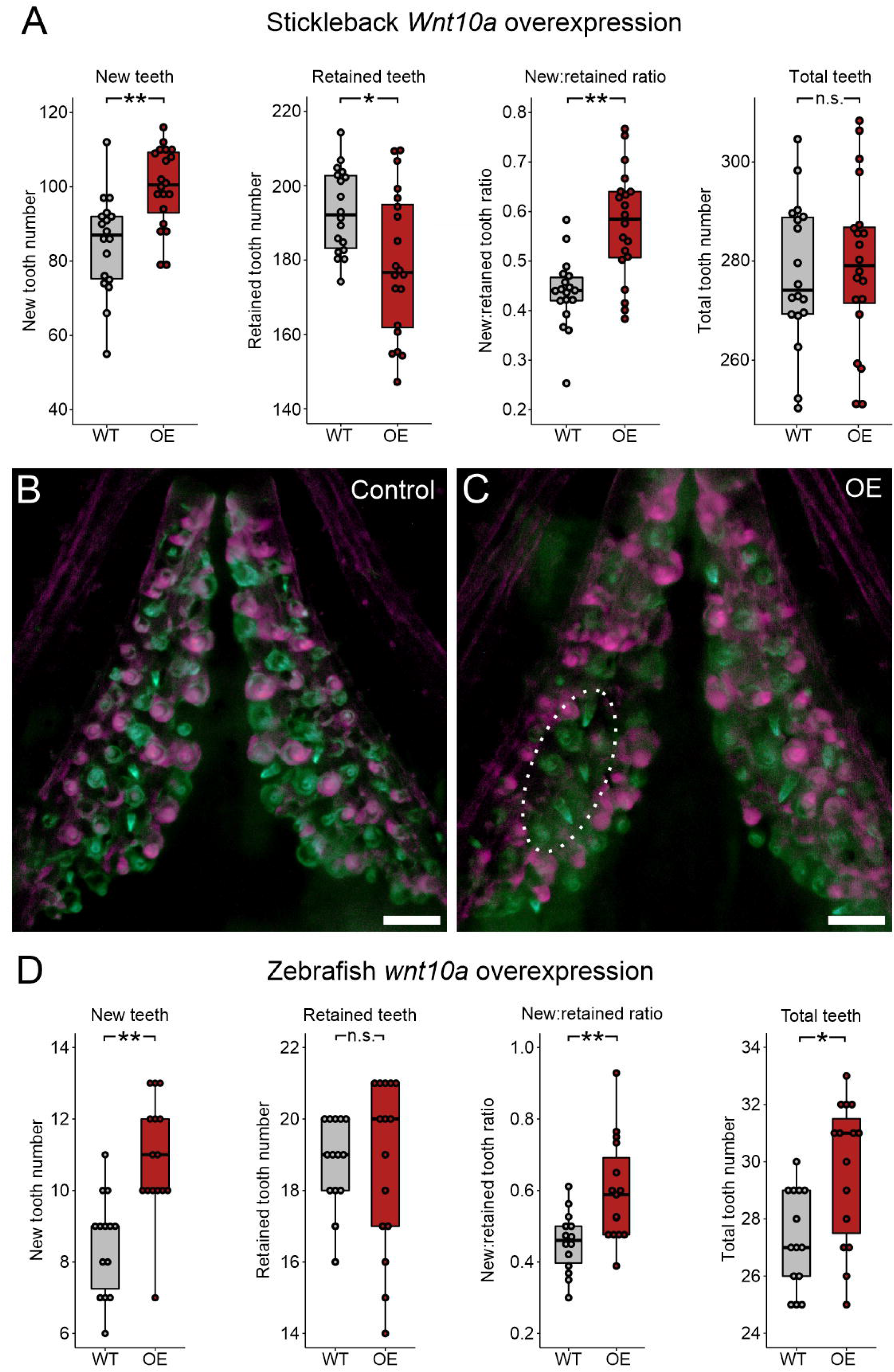
*Wnt10a* overexpression causes accelerated new tooth formation in stickleback and zebrafish. All *P-*values are derived from Wilcoxon Rank-Sum tests, adjusted for multiple hypothesis testing. **A.** In stickleback, the number of new teeth (*P*=0.0015), retained teeth (*P*=0.032), and the new:retained ratio (*P*=0.0012) showed significant differences between WT and OE fish, however the total number of teeth did not (*P*=0.77) (n = 18 control and 20 OE fish). **B,C.** Overlay images of stickleback ventral tooth plates showing Alizarin Red and calcein signal in control (B) and OE (C) individuals. Note clusters of new teeth (white dotted circle showing an example on the left VTP). **D.** In zebrafish, significant increases in the number of new teeth (*P*=0.0016), the new:retained ratio (*P*=0.0084), and total teeth (*P*=0.014) were found but retained tooth number (*P*=0.56) did not significantly change (n = 14 control and 15 OE fish). Boxes represent the 25th-75th percentiles, and the median is shown as a gray bar. Scale bars: 100 μm.

**Figure 4.**
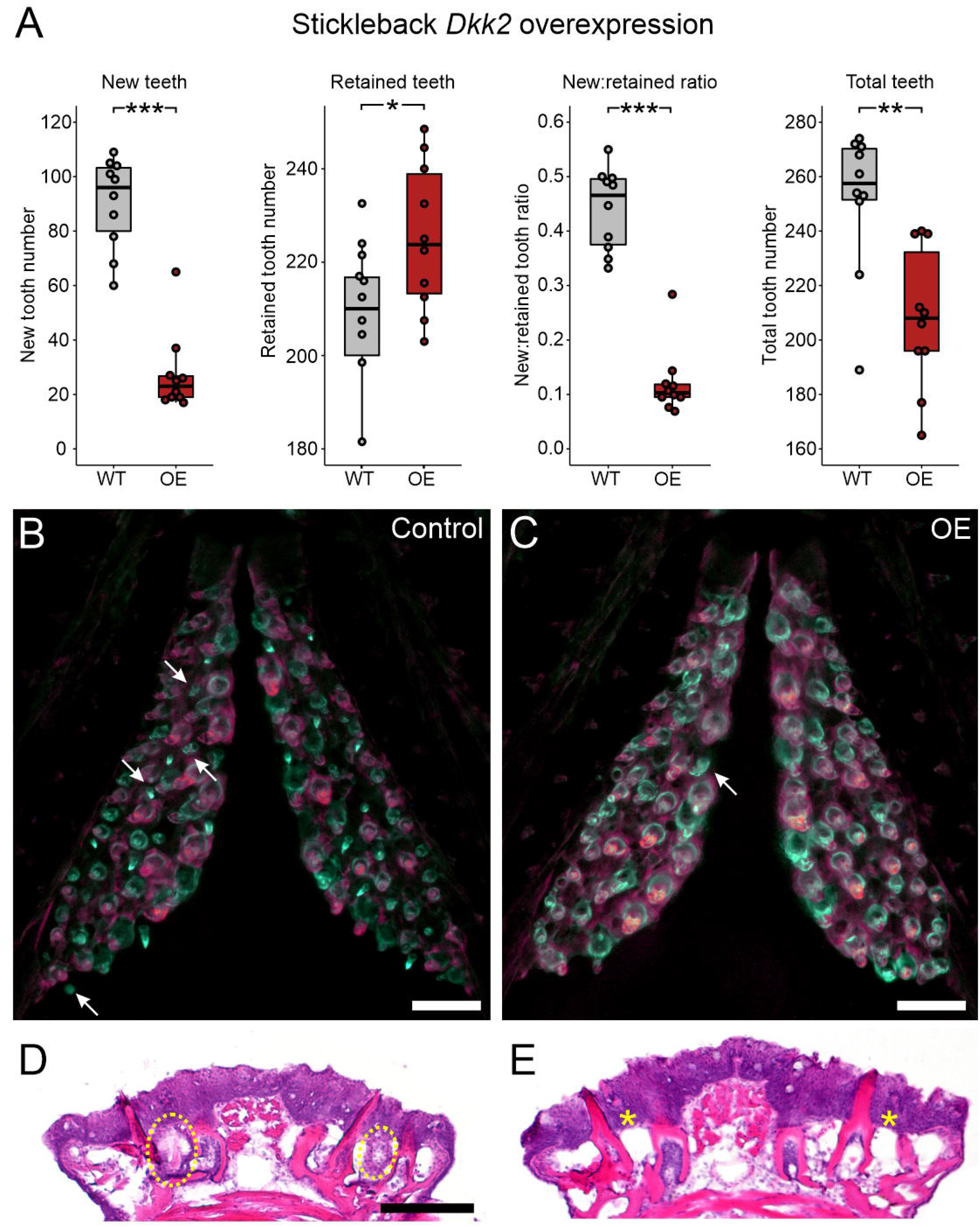
*Dkk2* overexpression diminishes new tooth initiation in sticklebacks. All *P-* values are derived from Wilcoxon Rank-Sum tests, adjusted for multiple hypothesis testing. **A.** Significant decreases in the number of new teeth (*P*=0.00049), the new:retained ratio (*P*=4.3e-5), and total teeth (*P*=0.0048), and an increase in retained teeth (*P*=0.037) were observed (n=10 control, 10 OE fish). **B,C.** Overlay images of stickleback ventral tooth plates showing Alizarin Red and calcein signal in control (B) and OE (C) fish. Note near complete absence of unankylosed new teeth in the OE fish (white arrow in C) compared to the WT fish (white arrows in B). The right side is unlabeled. **D,E.** Hematoxylin and Eosin (H&E) staining on transverse sections of tooth fields reveals no tooth germs of any stage in *Dkk2* OE fish (n=7/7), suggesting that this treatment does not cause tooth germs to arrest (yellow ovals in D show tooth germs, asterisks in E sit above positions normally populated by tooth germs). Boxes represent the 25th-75th percentiles, and the median is shown as a gray bar. Scale bars: 100 μm.

We next tested *wnt10a* OE in zebrafish (Fig. 3D) using the same 18-day pulse-chase overexpression interval, but with zebrafish resting and heat shock temperatures (see Methods and Fig. 2B). We found increased numbers of new teeth formed during the treatment interval under *wnt10a* OE. However, unlike sticklebacks, the number of retained teeth was not affected in zebrafish. Overall the new:retained tooth ratio was significantly increased. While we did not observe any change in the number of tooth families in any OE individual (n=15/15), we did find an increase in the total tooth number under *wnt10a* OE, due to a higher number of tooth families undergoing early replacement.

To address whether Wnt signaling inhibition could negatively influence tooth formation, we asked if Dkk2, a secreted Wnt signaling inhibitor, could decrease tooth replacement rates or total tooth number in sticklebacks (Fig. 4A). We found that *Dkk2* OE strongly reduced the presence of new teeth, while simultaneously increasing the presence of retained teeth. Together, these two effects led to a sharp decrease in the new:retained ratio, while also decreasing the number of total teeth. In the control condition, new teeth comprised mostly mid- or late bell stage tooth germs (Fig. 4B, arrows), whereas the few unankylosed new teeth we observed under *Dkk2* OE were always at or near the eruption stage (Fig. 4C, arrow). These results are generally reflected by each tooth field type alone (Fig. S5). To assess whether *Dkk2* OE could be stalling tooth germs at stages prior to bone deposition (at bud, cap, or early-bell stages), we analyzed tooth field histology using H&E-stained sections (Fig. 4D, E). We found no evidence of bud, cap, or early-bell stage tooth germs across any pharyngeal tooth field we observed from *Dkk2* OE individuals (n=7/7), suggesting that Dkk2 does not cause new or existing tooth germs to arrest during early differentiation. Measuring the area of the VTPs showed that tooth field area was not significantly altered after *Dkk2* OE (Fig. S3).

**Figure 5.**
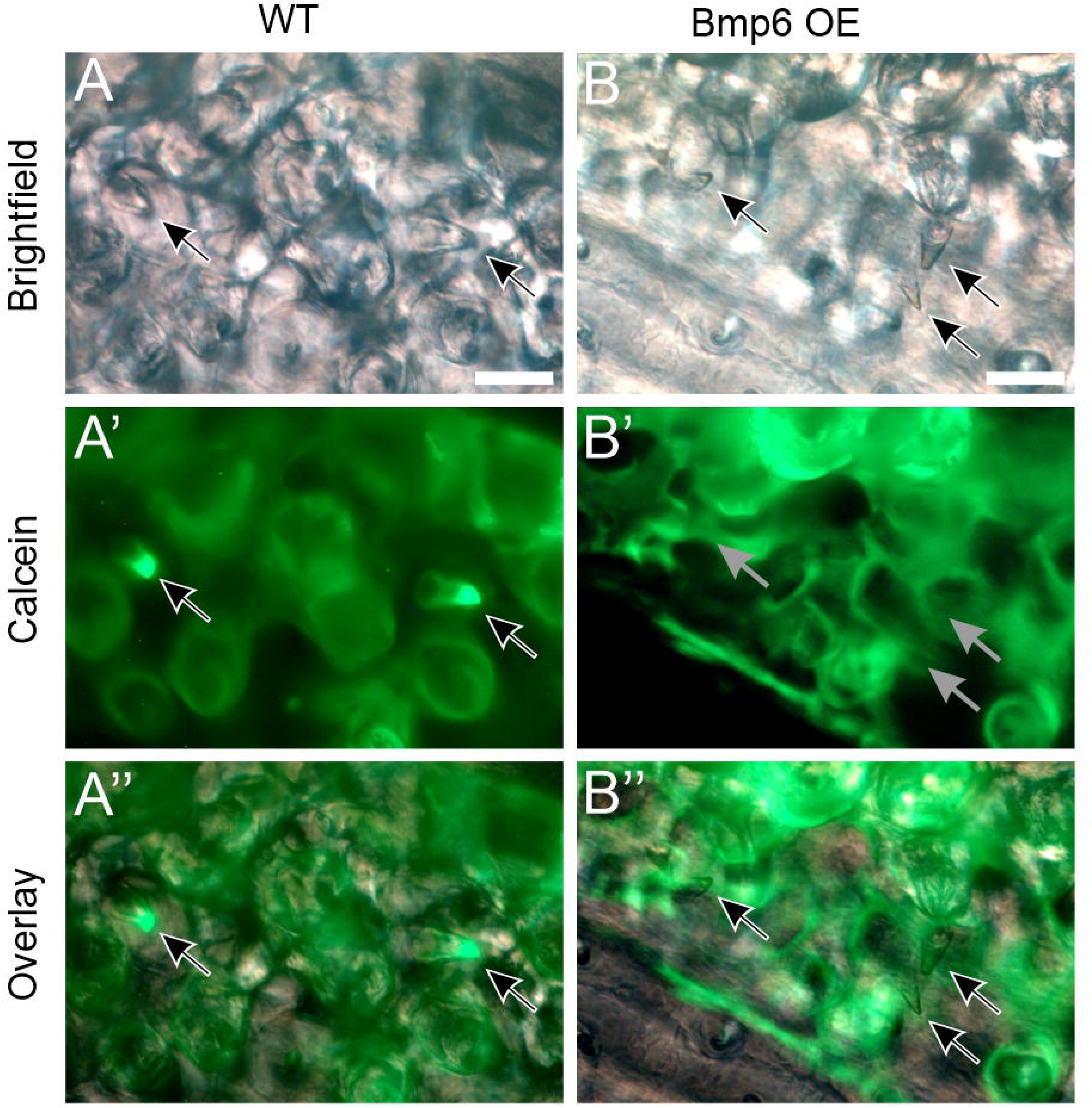
Stalled tooth germs resulting from *Bmp6* overexpression. A shows WT control, B shows OE transgene carrier. A and B show brightfield, A’ and B’ show green channel fluorescence (calcein), A’’ and B’’ show an overlay. Note that there was no caclein signal in the four tooth germs indicated in the *Bmp6* OE fish (gray arrows in B’). Black and white arrows otherwise mark tooth germs in those panels where they are visible. Scale bars: 25 μm.

### BMP pathway modulation by overexpression of *Bmp6* and *Grem2a*

We next sought to test whether modulation of BMP signaling could affect tooth replacement rates or total tooth number. Specifically, our hypothesis that a hair-like cycling mechanism operates in teeth predicts that the initiation of tooth replacement events should be inhibited by increased BMP signaling and promoted by BMP inhibition. To test this hypothesis, we conducted a *Bmp6* OE treatment, given known roles for *Bmp6* in epithelial appendage development and regeneration, including in stickleback teeth and mouse hair. *Bmp6* OE produced a striking tooth phenotype unique among the phenotypes observed in this study: late bell (bony) tooth germs that were negative for both Alizarin Red and calcein (Fig. 5). We interpret these unstained tooth germs as “stalled,” i.e. tooth germs that initiated bone growth just after the Alizarin pulse/treatment onset, but ceased differentiation and bone deposition during the treatment interval and were thus no longer producing bone by the end of the treatment and the calcein chase. We conservatively counted these unstained, stalled tooth germs as assumed “new” teeth for this treatment, which were unique to *Bmp6* OE and never otherwise observed. Even so, *Bmp6* OE resulted in sharply reduced new tooth formation (Fig. 6A). Surprisingly, *Bmp6* OE also resulted in a decrease in retained teeth, suggesting that Bmp6 negatively affects new tooth formation while also promoting the shedding of existing teeth. *Bmp6* OE overall led to a decrease in the new:retained tooth ratio, indicating that exogenous *Bmp6* inhibited tooth replacement rates overall by more strongly disrupting new tooth formation relative to its promotion of retained tooth loss. Given that we found reductions in both new and retained teeth, *Bmp6* OE necessarily decreased total tooth number, resulting in the treatment group having only ∼50-60% of the number of teeth that their WT control siblings possessed at the end of the OE treatment. Most of these trends are reflected by each tooth field type alone (Fig. S6). Qualitatively, we found large swaths of stickleback tooth fields that were devoid of erupted teeth (Fig. 6C, dotted oval), which was unique to this treatment condition. Measuring the area of the VTPs showed that tooth field area was not significantly altered under *Bmp6* OE (Fig. S3) despite the lower number of teeth present within each tooth plate. *Bmp6 OE* additionally caused body axis bending and novel rib-like bony protrusions to form on specific vertebral elements (caudal vertebrae 3-8; Fig. S7). This was the only overt morphological phenotype we observed arising from any heat shock experiment. We additionally performed a negative control assay on the *Bmp6* OE line, again allowing us to ascertain whether heterozygous transgene carriers in the absence of heat shocks exhibit alterations to any aspect of tooth turnover; as with the *Wnt10a* negative control experiment, we found no significant change in any measured variable (Fig. S4). Notably, we also found no calcein negative (stalled) tooth germs in this negative control experiment.

**Figure 6.**
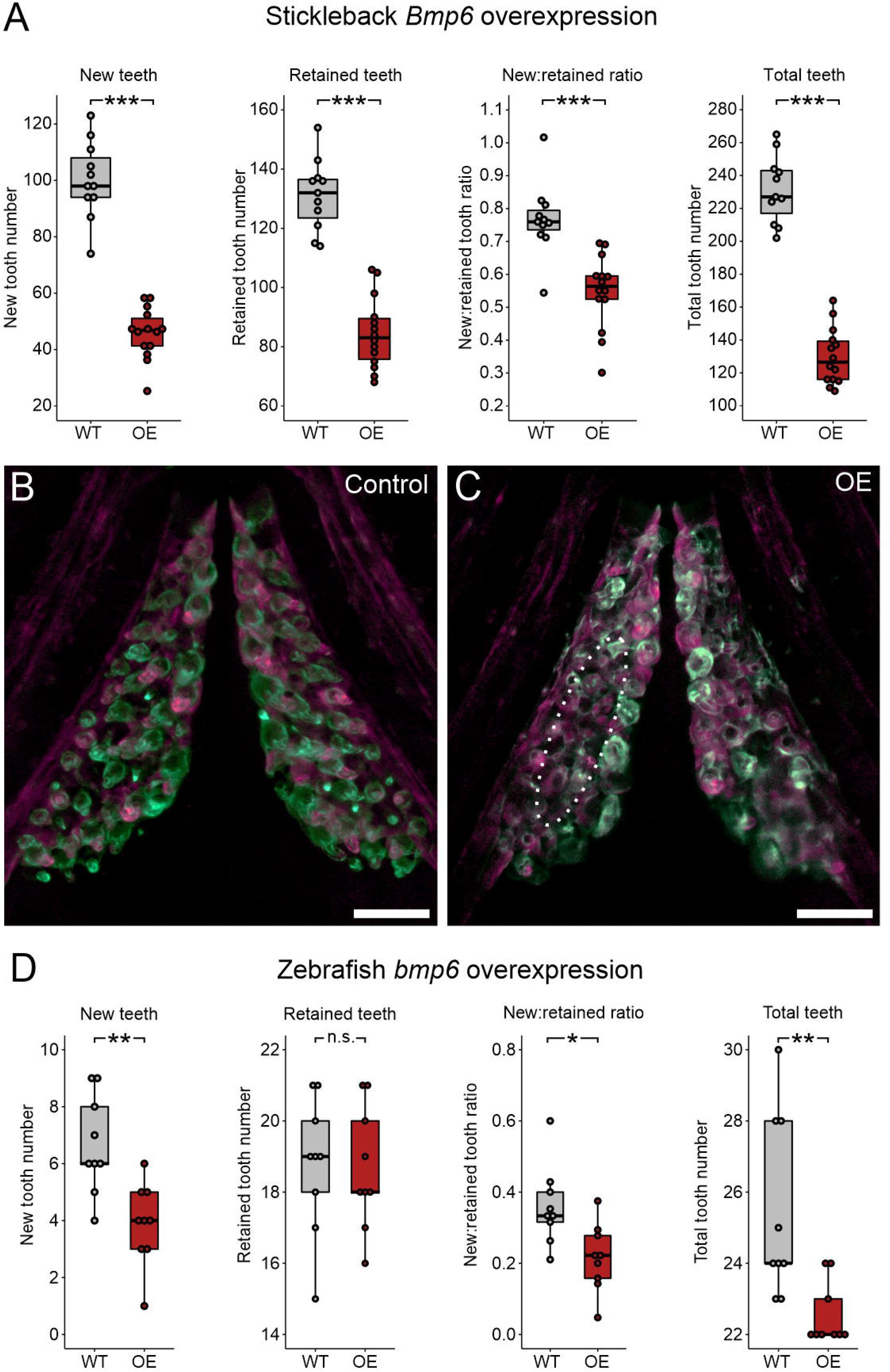
*Bmp6* overexpression limits new tooth development in stickleback and zebrafish. All *P-*values are derived from Wilcoxon Rank-Sum tests, adjusted for multiple hypothesis testing. **A.** In sticklebacks, significant decreases in all four variables were detected: the number of new teeth (*P*=3.7e-5), the number of retained teeth (*P*=3.7e-5), the new:retained ratio (*P*=4.4e-5), and total teeth (*P*=3.7e-5), (n = 11 control, 14 OE fish). **B,C.** Overlay images of stickleback ventral tooth plates showing Alizarin Red and calcein signal in control (B) and OE (C) fish. Note regions usually populated with ankylosed teeth are devoid of any such structure (dotted oval in D, right side is unlabeled). **D.** In zebrafish, significant decreases in the number of new teeth (*P*=0.0084), the new:retained ratio (*P*=0.018), and total teeth (*P*=0.0084) were detected but retained tooth number (*P*=0.79) did not significantly change (n = 9 control, 9 OE fish). Boxes represent the 25th-75th percentiles, and the median is shown as a gray bar. Scale bars: 100 μm.

**Figure 7.**
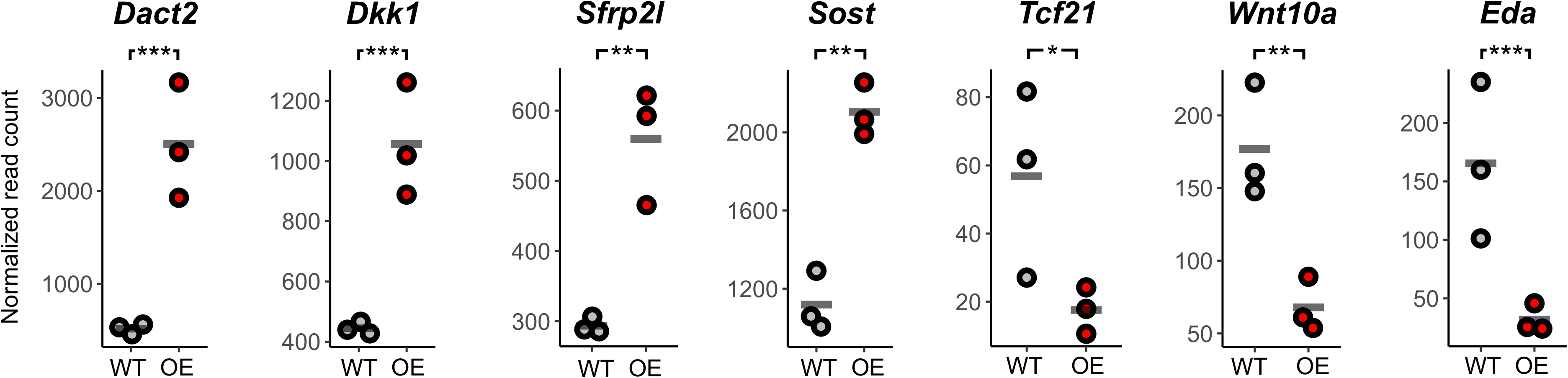
RNA-seq reveals transcriptional changes associated with *Bmp6* overexpression. N=3 each control and OE ventral tooth plates were collected ∼12 hours after a single heat shock and subjected to RNA-seq. Each fish’s normalized read count is shown for each gene, and a gray bar indicates the mean of each group. We found increased transcription of Wnt inhibitors (*Dact2*, *Dkk1*, *Sfrp2l*, and *Sost*) and reduced expression of *Wnt10a*, *Tcf21*, and *Eda*. \**P*<0.05, \*\**P*<0.01, \*\*\**P*<0.001 (Wald tests, corrected *P* for multiple hypothesis testing).

Bmp6 is thought to prevent hair regeneration initiation in part by inhibiting Wnt signaling (Kandyba et al., 2013; Wu et al., 2019). To ask whether *Bmp6* OE caused transcriptional changes in stickleback dental tissue that are consistent with Wnt pathway suppression, we performed RNA-seq on OE and control ventral tooth plates 12 hours after a single heat shock and compared their transcript profiles using DESeq2 (Love et al., 2014). *Bmp6* OE caused significant upregulations of multiple known Bmp effector genes, including multiple Smads (*Smad1*, *-2, -6a, -6b, -7, -9,* and *-10a*), *Msx1a*, *Msx2b*, and *Dlx3b*, as well as BMP inhibitors, including *Bambia, Grem1b, Grem2a, Nog1*, *Nog2*, and *Sostdc1a*. We additionally observed a significant increase in orthologs of Wnt inhibitor genes that have hypothesized roles in hair follicles, including *Dact2* (Meng et al., 2008)*, Dkk1* (Andl et al., 2002; Kwack et al., 2012; Plikus et al., 2011)*, Sfrp2l* (Kim and Yoon, 2014; Kim et al., 2010), and *Sost* (Avigad Laron et al., 2018) (Fig. 7). Alternatively, we found reduced transcription of *Wnt10a* and *Tcf21* (Fig. 7), however these were the only annotated Wnt or TCF/Lef genes with significantly reduced expression. We also found reduced expression of *Ectodysplasin*, a gene with known positive roles in tooth and hair differentiation (Aigler et al., 2014; Harris et al., 2008; Jandzik and Stock, 2021). Together, these data suggest that exogenous BMP signaling via *Bmp6* OE is sufficient to cause transcriptional changes consistent with the inhibition of Wnt signaling, as expected from a hair regeneration model.

Given the observed upregulation of Wnt inhibitors and downregulation of *Wnt10a*, we next asked whether sustained *Bmp6* OE was sufficient to alter the spatial distribution of TCF/Lef transcription factor activity. We thus performed a 10 heat shock *Bmp6* OE experiment on fish that carried a previously described TCF/Lef synthetic reporter construct (Shimizu et al., 2012; Square et al., 2021). In stickleback tooth fields, this reporter is active in the inner and outer dental epithelium of all tooth germs as well as the Successional Dental Epithelium (SDE) (Fig. 8A,B) i.e. the hypothesized precursor tissue of tooth germ epithelia that demonstrates a gene expression signature similar to the zebrafish successional dental lamina (Square et al., 2021). We found that *Bmp6* OE did not visibly reduce TCF/Lef reporter activity in developing tooth germs, which appeared unaltered in their reporter expression (Fig. 8, arrows). Conversely, we observed a significant reduction in the number of reporter-positive SDE in *Bmp6* OE, with 4/9 treatment individuals demonstrating zero TCF/Lef reporter-positive SDE at any tooth field (Fig. 8B-D). This result suggests that *Bmp6* OE works to inhibit Wnt signaling specifically in naïve dental epithelia (the SDE).

**Figure 8.**
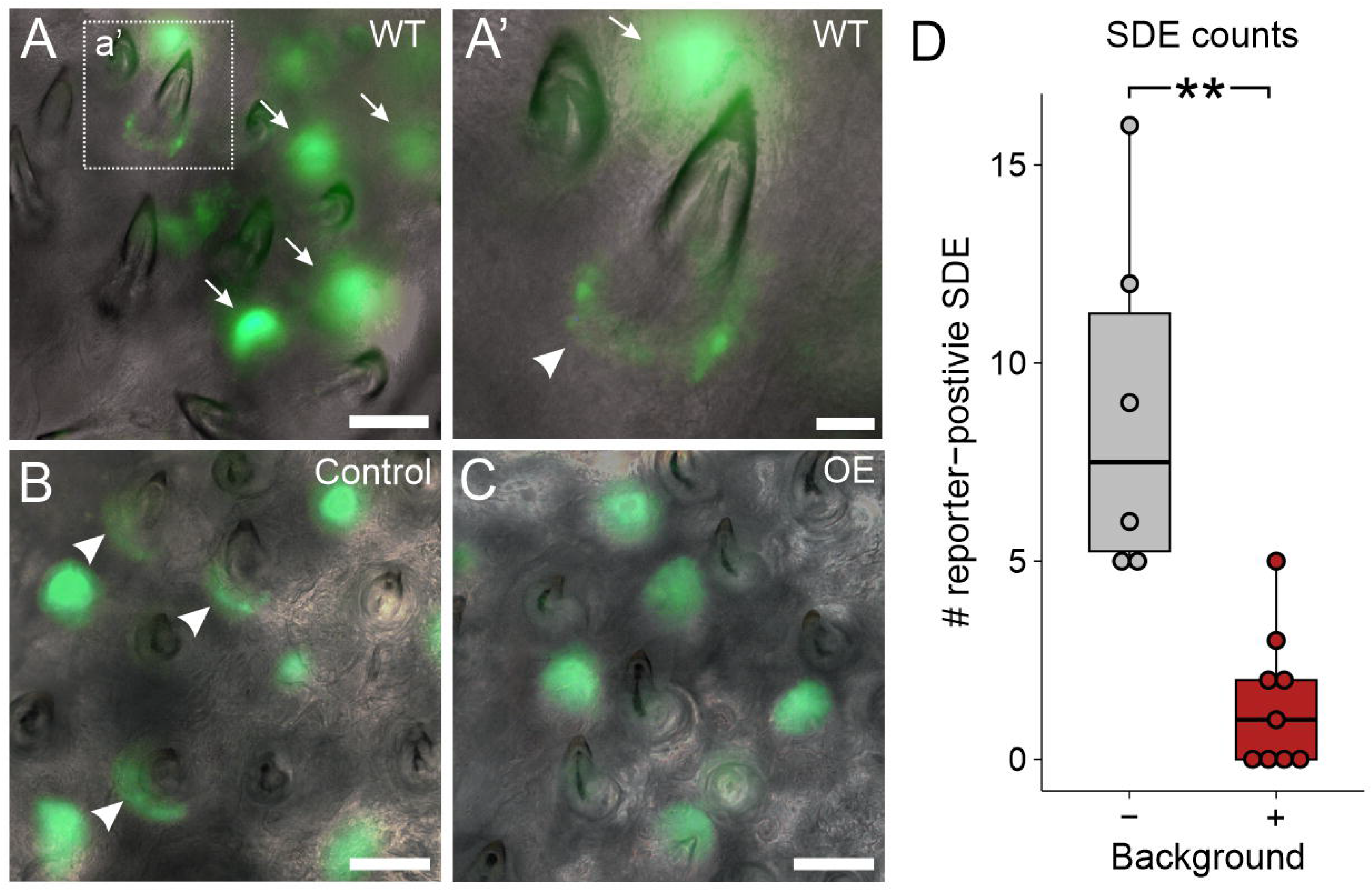
*Bmp6* OE caused a reduction of TCF/Lef reporter activity in successional dental epithelia (SDE). A,B. WT images. Arrows mark tooth germs, arrowheads mark SDE. **C,D** control and OE examples, respectively. Note the GFP-positive SDE in the control individual (arrowheads). **E.** A box and whisker plot showing the number of TCF/Lef-positive SDE. Wilcoxon Rank-Sum test *P*=0.0021 (n=6 control, 9 OE fish). Scale bars: 100 μm in A-C, 25 μm in A’.

We next asked if *bmp6* OE had any of these same effects on zebrafish tooth fields (Fig. 6D). While we did not observe any stalled tooth germs as we did in sticklebacks, we did find a significant decrease in the number of new teeth that formed during the treatment interval, consistent with a negative role in tooth germ initiation and/or differentiation. Unlike sticklebacks, zebrafish exhibited no significant differences in retained tooth number under *bmp6* OE, while the new:retained ratio fell significantly. We did not observe any changes to the number of tooth families present (n=9/9), however we did detect a significant decrease in the overall number of teeth, likely brought on by the paucity of new bell stage tooth germs in the OE condition.

To test whether BMP pathway inhibition could promote tooth turnover, we performed an overexpression experiment using the predicted BMP antagonist *Grem2a* in sticklebacks (Fig. 9A). *Grem2a* OE did not significantly change new tooth number but did decrease retained tooth number. This drop in the retained tooth number per fish was sufficient to increase the new:retained tooth ratio, achieving a higher relative proportion of new teeth by eliminating retained teeth. *Grem2a* OE also significantly reduced total tooth number. Most individual tooth field types had such trends (Fig. S8). Measuring tooth field area showed no significant alteration under *Grem2a* OE (Fig. S3).

**Figure 9.**
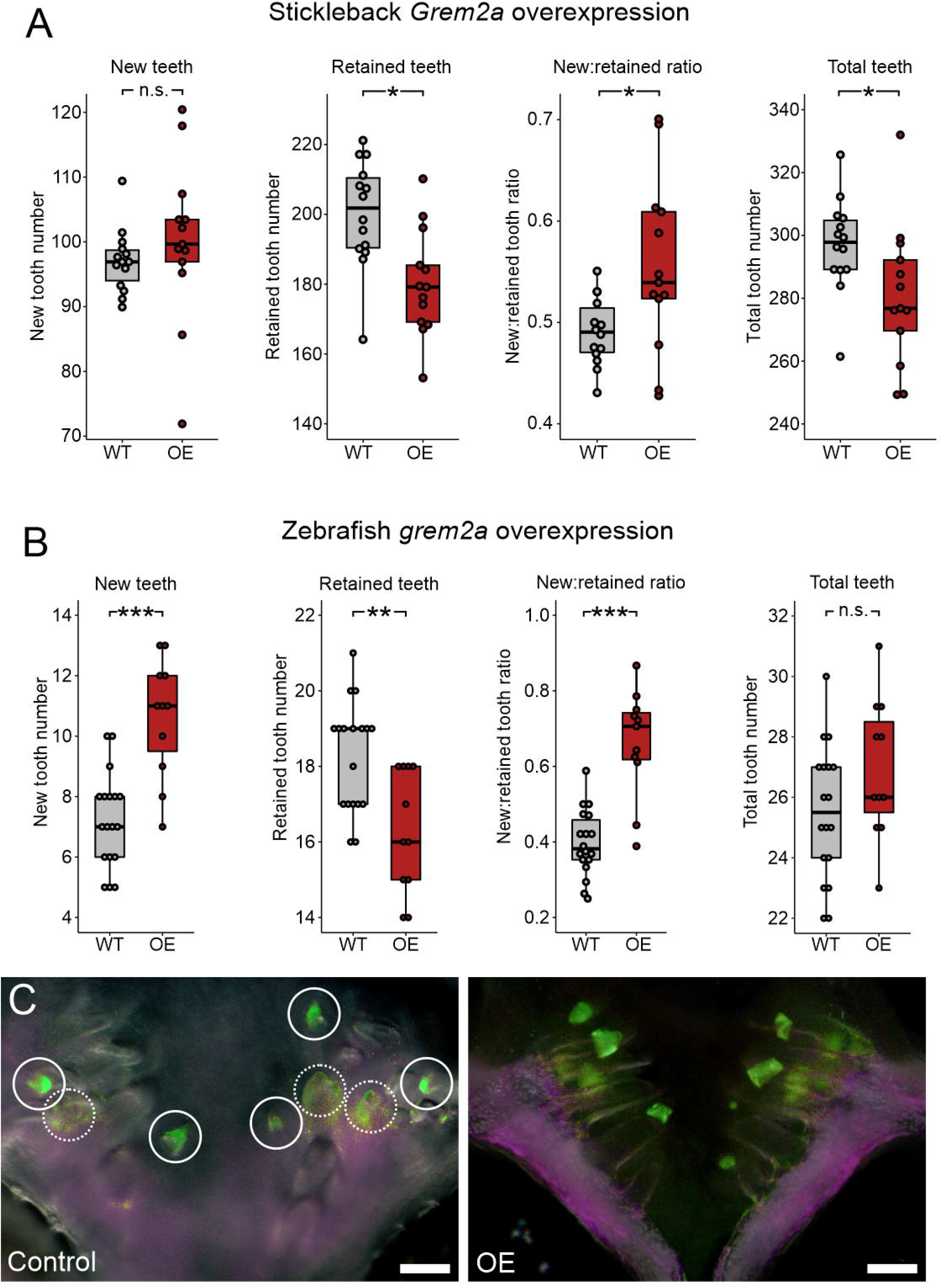
Effects of *Grem2a* overexpression in stickleback and zebrafish. All *P-*values are derived from Wilcoxon Rank-Sum tests, adjusted for multiple hypothesis testing. **A.** In sticklebacks, significant decrease in the number of retained teeth (*P*=0.018), a significant increase in the new:retained ratio (*P*=0.039), and total teeth (*P*=0.029), but no significant changes to new teeth (*P*=0.11) were observed (n=14 control, 13 OE fish). **B.** In zebrafish, significant increases in the number of new teeth (*P=*0.00062) and the new:retained ratio (*P=*0.00053), a significant decrease in retained teeth (*P*=0.0062), and no significant change in total teeth (*P*=0.14) was observed (n=18 control, 11 OE fish). **C,D.** Overlay images of zebrafish ventral tooth plates showing Alizarin Red and calcein signal in control (B) and OE (C) fish. White circles indicate new teeth (solid circles mark superficial teeth, and dotted circles mark teeth deep in the tooth field that are occluded by erupted teeth). Boxes represent the 25th-75th percentiles, and the median is shown as a gray bar. Scale bars: 100 μm.

Given that *Grem2a* likely inhibits a subset of BMP ligands, we sought to test whether broader BMP pathway inhibition would cause similar effects on dental phenotypes. We thus performed pulse-chase bone labeling on wild type sticklebacks that were treated with either DMSO or LDN 193189 (“LDN"), a selective inhibitor of Bmpr1a/Alk3 and Acvr1/Alk2 signal transduction. We found that, like in the *Grem2a* OE experiment, the number of retained teeth significantly decreased under LDN treatment, while the number of new teeth did not significantly change (Fig. S9). Together, these differences drove an increase in the new:retained tooth ratio, and a decrease in the total number of teeth (Fig. S9), again mirroring the *Grem2a* OE result.

We next tested *grem2a* OE in zebrafish to ask whether BMP signaling inhibition also promoted tooth replacement in this species (Fig. 9B-D). This treatment resulted in an increase in the new:retained tooth ratio, which arose from a simultaneous increase of new teeth and a reduction in retained teeth. Zebrafish *grem2a* OE did not alter the number of tooth families observed (n=11/11), nor did it cause a significant change in the total number of teeth.

## Discussion

### Endogenous expression domains of stickleback *Wnt10a, Dkk2, Bmp6*, and *Grem2a* reflect similar germ layer partitioning as in hair follicle gene expression

As a hair regeneration model would predict, *Wnt10a*, *Dkk2*, *Bmp6*, and *Grem2a* exhibit expression in tooth organ epithelium and mesenchyme that appears cyclic during tooth replacement. These four genes additionally show expression partitioning between epithelial and mesenchymal germ layers in ways that overall resemble previously reported expression domains of the mammalian orthologs of these genes in hair follicles (detailed below). Overall, we found that the dental mesenchyme and inner dental epithelium of bell stage tooth germs expresses all four of these secreted ligands. We additionally found expression of all four genes surrounding tooth germs within the tooth field (brackets throughout Fig. 1). The apparently high representation of different secreted ligands suggests that tooth fields are subject to a rich and dynamic amalgam of Wnt and BMP signaling molecules overall. The endogenous expression patterns of these secreted factors in tooth fields thus leaves open the possibility that they participate in some form of reaction-diffusion system between tooth organs and/or more broadly as parts of regulatory centers or waves that coordinate tooth replacement; such regulatory systems have been hypothesized to explain different coordinated aspects of hair field growth or regeneration (Plikus et al., 2008; Sick et al., 2006; Plikus et al., 2009; Chen et al., 2014; Nagorcka, 1983).

*Wnt10a* was focally expressed in both epithelium and mesenchyme of developing tooth germs, but epithelial expression appeared to wane by late bell stages, and mesenchymal expression was essentially absent in erupted tooth mesenchyme (Fig. 1A-D). This expression roughly corresponds to the cycle previously observed in the mouse hair follicle, where *Wnt10a* expression was found in the early anagen (growth) phase, but not telogen (the quiescent resting phase which ends with hair shedding) (Reddy et al., 2001). The Wnt inhibitor *Dkk2* is thought to confer a feedback response in mouse hair follicles, wherein its expression in hair follicle mesenchyme (the dermal papilla) is brought on by Wnt effectors and helps trigger the end of the growth (anagen) phase (Harshuk-Shabso et al., 2020). Stickleback mesenchymal expression domains of *Wnt10a* and *Dkk2* are generally consistent with such a relationship, showing temporal complementarity: *Wnt10a* shows marked expression at the beginning of tooth differentiation (bud stage) that wanes throughout differentiation (Fig. 1A-D), whereas *Dkk2* appears to be excluded from bud stage mesenchymal cells, transcribed during bell stages, and is then highly expressed during and after eruption, especially in fully differentiated tooth mesenchyme (odontoblasts; Fig. 1E,F).

BMP pathway genes additionally show expression domains in stickleback tooth fields that are partitioned by germ layer and stage in a way that roughly corresponds to mouse hair follicles. *Bmp6* is highly expressed during growth (anagen) stages of the mouse hair follicle cycle (Wu et al., 2019), similar to the inner dental epithelial and mesenchymal expression that is present during early tooth germ growth (Cleves et al., 2014; Square et al., 2021) (Fig. 1H). Despite this marked expression of *Bmp6* during organ differentiation, *Bmp6* has been shown to delay organ initiation in hair follicles by inhibiting the telogen to anagen transition (Wu et al., 2019). Broad similarities exist with teeth: in both stickleback and zebrafish, *Bmp6* demonstrates punctuated upregulation in specific dental tissues during tooth organ differentiation (Square et al., 2021) (Fig. 1G-J), while also preventing the formation of new tooth organs (Fig. 6A,D). *Grem2* expression has not been analyzed in mouse hair to our knowledge, but *GREM2* expression in human hair follicles is found in the mesenchymal dermal sheath cup surrounding the hair follicle, which has led to hypotheses that it regulates the hair follicle stem cell niche (Niiyama et al., 2018; Niiyama et al., 2022). Like human *GREM2* in hair fields, stickleback *Grem2a* expression in dental fields was found most strongly in mesenchyme surrounding tooth germs (Fig. 1K,M, brackets), though was generally more widespread.

### Opposing roles of the Wnt and BMP pathways in epithelial appendage regeneration

We parsimoniously hypothesized that different epithelial appendages, including teeth, could use a shared input scheme to dictate the progression of regeneration. To test the hypothesis that teeth and hair share instructive functions for secreted ligands during regeneration, we used a candidate gene approach to ask whether specific Wnt and BMP pathway members could regulate tooth replacement rates in a manner consistent with their known or suspected roles in the Wnt-BMP mechanism described in hair follicles. Our hypothesis specifically predicted that Wnt pathway stimulation or BMP pathway inhibition would increase the replacement rate of teeth, while BMP pathway stimulation or Wnt pathway inhibition would inhibit regeneration. Overall, our results supported our hypothesis: we found that *Wnt10a* or *Grem2a* OE increased tooth replacement rates (as measured by the new:retained tooth ratio) in both sticklebacks and zebrafish, while *Bmp6* or *Dkk2* OE both decreased the initiation of new teeth, the former in both sticklebacks and zebrafish, the latter in sticklebacks only (we did not test zebrafish *dkk2* OE). Our interpretation of these results is that tooth regeneration is indeed modulated in a congruous fashion as hair with respect to the known opposing roles of the Wnt and BMP pathways. Combined with a previous study showing that a common genetic battery marks hair follicle stem cells and naïve successional dental epithelia (SDE) (Square et al., 2021), these data support a model where teeth and hair, as epithelial appendages, share evolutionarily conserved genetic instructions used to regulate whole organ regeneration. Using RNA-seq, we also show that stickleback orthologs of genes hypothesized to be involved in maintaining mouse hair follicle stem cell quiescence, like *Dkk1*, *Sfrp2*, and *Sost*, are specifically upregulated under stickleback *Bmp6* OE in dental tissue, while *Wnt10a* transcription is suppressed (Fig. 7). Furthermore, based on TCF/Lef reporter activity (Fig. 8), *Bmp6* appears to negatively affect the canonical Wnt pathway in the SDE (arrowheads in Fig. 7), but not in differentiating teeth (arrows in Fig. 7), providing further support for a specific role for *Bmp6* in the activation of epithelial organ differentiation.

### Regeneration rate and total tooth number can be influenced separately or in concert

Zebrafish typically end primary expansion of their single pair of tooth fields at around 30 days old, forming a consistent arrangement of 11 tooth families per ventral pharyngeal tooth field that is maintained into adulthood (Van der Heyden and Huysseune, 2000). In all three of our zebrafish OE experiments, the replacement rate was significantly altered, but the number of tooth families was not altered. Thus, the differences in zebrafish total tooth number we documented (which includes tooth germs) were likely reflecting the higher (*wnt10a* OE) or lower (*bmp6*) number of tooth families that were actively undergoing regeneration, causing there to be two ossified teeth at such tooth positions. Conversely, zebrafish *grem2a* OE increased the replacement rate by both increasing new teeth and decreasing retained teeth, without changing the number of tooth families or significantly increasing total tooth number. Overall, these data suggest that zebrafish, like sticklebacks, can adjust tooth replacement rates without demonstrating significant changes to tooth family number or total tooth number.

Secreted signals that simply promote or inhibit the differentiation of all tooth germs – replacement and primary – would be predicted to change the replacement rate while also changing total tooth number in the same direction in concert. We found that *Dkk2* OE in sticklebacks best fit the predicted phenotype arising from an exogenous secreted protein that inhibits all tooth formation: new teeth, the replacement rate, and total teeth all dropped under *Dkk2* OE, while retained teeth rose, indicating that tooth turnover and primary growth were both precipitously slowed (Fig. 4A). Notably, *Dkk2* OE appears to have allowed bell stage tooth germs that were present at the time of OE onset to finish development (since zero early, mid- or late bell stage tooth germs were observed in OE fish).

Both *Bmp6* and *Grem2a* OE treatments in sticklebacks caused a drop in retained tooth number that was not accompanied by an increase in new tooth number. These results strongly suggest that tooth shedding events can be promoted in the absence of increased new tooth formation in sticklebacks. Particularly in the *Bmp6* treatment, tooth shedding increased in tandem with a severe reduction in the number of new teeth (Fig. 6A). Thus, in sticklebacks, the process of tooth replacement can apparently be regulated not just by influencing the start of replacement tooth formation, but also by independently regulating tooth shedding. These data further support the hypothesis (Ellis et al., 2015; Square et al., 2021) that the documented evolutionary changes in stickleback tooth number and replacement rates could manifest by changes to tooth shedding mechanics separately from the tooth formation process.

### Other differences and similarities between sticklebacks and zebrafish

Each of our OE treatments affected at least one aspect of tooth replacement in a direction predicted by a hair regeneration model. However, the specific nature of each response was not always the same between sticklebacks and zebrafish for the three gene ortholog pairs we tested in both species (*Wnt10*, *Bmp6*, and *Grem2a*). Importantly, the lack of an effect in one species versus the other could simply reflect limitations of our OE assay: for example, the genomic integration of the OE construct in one species could feasibly be at a genomic location that allows for stronger or weaker transgene expression upon heat shock, causing us to observe unique effects. It is also possible that species-specific protein sequence differences contribute to the differences we observed. Despite these limitations of our approach, we are still able to deduce important similarities and likely differences in how these two fishes respond to the overexpression of Wnt and BMP pathway members.

The number of new teeth was increased by *Wnt10a* OE and decreased by *Bmp6* OE in both fish species, suggesting that these secreted factors exert opposite, conserved effects on the initiation of replacement tooth growth. While *Grem2a* did work to increase the new:retained ratio in both species, this was achieved by unique means: in sticklebacks, by only reducing the number of retained teeth (Fig. 9A), and in zebrafish by simultaneously reducing the number of retained teeth and increasing the number of new teeth (Fig. 9B). *Grem2a* thus may exert some effects in zebrafish that are not realized in sticklebacks.

In zebrafish, significant changes in the tooth replacement rate (as measured by the new:retained ratio) were always at least partly driven by corresponding changes to the number of new teeth. Furthermore, zebrafish retained tooth numbers showed either no detectable change (*wnt10a* [Fig. 3D] and *bmp6* [Fig. 6D]) or changed in the opposite direction as new teeth (*grem2a*; Fig. 9B). Conversely, sticklebacks showed changes in the new:retained tooth ratio that were driven either just by changes in retained tooth number (*Grem2a*; Fig. 9A) or where both retained teeth and new teeth changed in the same direction (both decreased under *Bmp6;* Fig. 6A). Thus, sticklebacks twice exhibited the ability to change retained tooth number in a manner that was unlikely a response to the increased generation of new teeth (because new teeth did not increase), whereas zebrafish did not demonstrate this phenomenon. These differences in response between sticklebacks and zebrafish could be partly due to the differences in regeneration strategy exhibited by these fish species: zebrafish adults maintain a set number (11) of stationary tooth families per tooth field that undergo one-for-one replacement in morphologically separated cycles, whereas sticklebacks have hundreds of teeth that do not retain a consistent arrangement into adulthood and appear to occasionally engage in one-for-two tooth replacement events (Square et al., 2021). We speculate that the highly canalized tooth regeneration process in zebrafish might contribute to less flexibility in the timing of tooth shedding during replacement tooth growth than in sticklebacks.

## Conclusions

The work presented here provides baseline evidence that tooth replacement can be governed by specific gene orthologs that have been shown to participate in hair organ regeneration and replacement in mammals (i.e. the Wnt-BMP oppositional mechanism). We additionally showed that, in sticklebacks, *Bmp6* promoted the expression of key Wnt inhibitors, like *Dkk1*, *Sfrp2l*, and *Sost*, whose orthologs are specifically implicated in mouse hair regeneration. These data overall suggest that epithelial appendages could share deeply homologous genetic architecture that drives basic developmental processes, despite the starkly dissimilar attributes of the functional organs. Our data additionally highlight previously unknown flexibility in the regulation of tooth replacement: it is possible to significantly speed up or slow down the tooth replacement cycle in both fish species tested here. Sticklebacks also demonstrated the ability to increase tooth shedding in the absence of additional new teeth, suggesting that active tooth shedding need not be caused by dissociation brought on by either a replacement tooth germ or damage/wear. Together, these results shed light on important regulatory mechanisms that affect tooth development and regeneration.

## Methods

### Overexpression transgene constructs

We used restriction-ligation cloning per standard methods to create the heat-shock overexpression construct plasmid backbone used in all seven OE treatments described here. We digested the pT2He-eGFP plasmid (Howes et al., 2017) with SfiI and BglII, discarded the smaller insert, and ligated the annealed oligos AGGCCCCTAAGGACTAGTCATATGTCTAGACTCGAGCCTAGGGGCGCGCCGGATCCA and GATCTGGATCCGGCGCGCCCCTAGGCTCGAGTCTAGACATATGACTAGTCCTTAGGGGCC TATC onto the ends. This created a plasmid with a multiple cloning site between the forward and reverse Tol2 transposase recognition sequences, including AscI, AvaI, AvrII, BamHI, BglII DraII, SpeI, XbaI, and XhoI restriction enzyme cut sites. We named this intermediate plasmid “T2Rv10.” Next, we added the SV40 poly adenylation signal by amplifying it from the pT2He-eGFP reporter construct with the primers GCCGAGATCTCGATGATCCAGACATGATAAG and GTTGTTGAATTCCCATACCACATTTGTAGAG and ligated into T2Rv10 via restriction cloning with BglII and EcoRI. Thereafter we used the primers ATAGGCCAGATAGGCCTCAGGGGTGTCGCTTGG and AATTGACTAGTCTTGTACAGCTCGTCCATGC to amplify the zebrafish *hsp70l* promoter and the *mCherry* coding sequence (without the stop codon) in tandem from the pT2He-mCherry reporter construct and ligated the product into the plasmid via the SfiI and SpeI restriction sites. Finally, we used the primers AATTGACTAGTGGCAGCGGTGCCACC and GCCGTCTAGAGGGTCCGGGGTTCTCTTC to amplify the Porcine Teschovirus-1 2A (P2A) self-cleaving peptide coding sequence from the pMS48 plasmid and ligated it into the plasmid via restriction cloning with SpeI and XbaI, yielding the “pT2overCherry” construct used in this work. This plasmid was thereafter outfitted with coding sequences of interest (with a stop codon) downstream of the P2A coding region using standard restriction site cloning with any two of the AscI, AvrII, BamHI, BglII, XbaI, or XhoI recognition sites that remain available in the multiple cloning site.

Seven coding regions for gene overexpression were synthesized by Gene Universal (Delaware, USA): stickleback *Wnt10a*, *Dkk2, Bmp6*, and *Grem2a*, and zebrafish *wnt10a*, *bmp6*, and *grem2a* (see Table S1 for accession numbers and full DNA sequences). These products were synthesized and cloned into pBlueScript SkII+ by Gene Universal, using XbaI and XhoI restriction enzyme recognition sites in all cases save stickleback *Wnt10a*, for which XbaI and BamHI were used due to an internal cut site for XhoI. Upon receiving these synthesized products, we digested the plasmids using the same restriction enzyme pair that was used to place them into pBlueScript and ligated them into pT2overCherry (digested with the corresponding restriction enzymes and the small insert removed). Ligation products were transformed per standard methods. Colony PCR screening was then performed to identify colonies carrying likely successful ligation products. These were miniprepped (Qiagen) and their full inserts were Sanger sequence verified, leaving ∼1 mL of bacterial culture at 4° C to later inoculate a larger liquid culture for midiprep. Once verified, midipreps (Qiagen), phenol:chloroform extractions, DNA precipitation, and resuspension in DEPC-treated water was performed per standard methods to prepare plasmids for injection.

### Fish husbandry and transgenic line establishment

Zebrafish and stickleback broodstock were raised and maintained under standard conditions. Transgenesis was accomplished by injecting Tol2 mRNA and pT2overCherry plasmids containing the aforementioned coding regions (see Table S1). F0 injected fish were outcrossed to WTs, founders were identified, and a single F1 offspring was thereafter used to establish a stable line from each founder. If the F2 generation exhibited significantly more than 50% transgenic offspring (Fisher’s Exact test), we outcrossed the line to WT until we observed ∼50% transgenic offspring (eliminating insertions until we inferred there was only a single insertion). The work presented here makes use of a single insertion of each transgene for each OE treatment.

### Heat shock treatments and pulse-chase bone labeling

To initiate an OE experiment, groups of ∼15-20 sibling fish were selected for treatment based on transgene carrier status. Such sample sizes were selected such that we could reliably detect an effect size of 20% or greater. We inferred transgene carrier status by lightly sedating fish in 50 mg/L MS-222 and briefly observing their lenses under red channel fluorescence, where the zebrafish *hsp70l* heat shock promoter drives sustained mCherry expression in the absence of heat shock. For a given experiment, WT and OE sibling fish were always raised together in a common tank for their entire lives and were only separated briefly from each other during the aforementioned sorting process. To initiate the treatment, groups of fish were placed into a tank containing a previously described (Ellis et al., 2015) Alizarin Red live-staining solution (0.1 g/L Alizarin Red S with 1mM HEPES) made using either stickleback or zebrafish tank/system water for each species. Sticklebacks were pulsed with Alizarin Red for 24-30 hours in 2L of solution, zebrafish were pulsed for 48-54 hours in 1L of solution. After the Alizarin Red staining pulse was completed, fish were rinsed once then washed 3 times for 10-30 minutes each in fresh tank/system water before being placed into the heat shock tank. Stickleback heat shock tanks were 4L in volume, were lightly aerated with an air stone for increased water agitation (to more uniformly distribute the temperature throughout the tank) and featured two 50-watt aquarium heaters set to 29° Celsius controlled by a timer. The timer engaged the heaters twice per day for two hours per pulse, starting every 12 hours. Since it takes the 4L tanks approximately 50 minutes to ramp up to the heat shock temperature, the two hours of applied heat translates to about 70 minutes at the heat shock temperature threshold (Metzger et al., 2016) per heat shock. Water changes were performed as needed every ∼5 days, avoiding the heat shock intervals. Zebrafish heat shock tanks essentially followed a published protocol (Duszynski et al., 2011), with some modification. Standard 2.8L Aquaneering tanks with a single 50W tank heater set to 39° C were kept on either a dripping water flow rate or with no flow or aeration (normal zebrafish movement was sufficient to agitate the water and create a uniform temperature throughout the tank). The timer activated the heater twice per day for 90 minutes, starting every 12 hours. Since it takes ∼2.8L of zebrafish system water about 20 minutes to heat to 39° C, these treatments brought fish to the heat shock temperature for around 70 minutes per treatment, as in the stickleback heat shock treatments. Water changes were performed at least once every four days by turning the tank flow on high for 10+ minutes, avoiding the heat shock intervals. After the 36^th^ heat shock on the 18^th^ day of the treatment, both species of fish were withheld from feeding, removed from their respective heat shock tanks, and stained in a calcein live staining solution (Ellis et al., 2015) (0.05 g/L calcein with 1mM sodium phosphate) for 16-18 hours, in 2L for sticklebacks and 1L for zebrafish. After rinsing and washing the fish in their corresponding tank/system water at least three times over 30 min, fish were euthanized in 250 mg/L MS-222, sorted by red channel fluorescence for gene overexpression activity (mCherry), and fixed in 4% PFA overnight at 4° C with high agitation.

### Negative control pulse-chase bone labeling assays

To test whether transgene carriers demonstrated differences in their new, retained, or total tooth numbers, we performed negative control pulse-chase assays using the same staining interval (18 days) but with no intervening heat shocks. Using both the *Wnt10a* and *Bmp6* OE transgenes, we repeated the pulse-chase bone labeling assay, except the heaters were removed from the 4L “treatment” tank, instead keeping them at normal rearing temperatures (∼18° C). Notably, these fish were full siblings to the treatment fish described in the results section, helping to control for familial genetic variation that may influence tooth replacement dynamics. We found no significant differences in the number of new teeth, number of retained teeth, the new:retained tooth ratio, or the total tooth number in either negative control experiment (Fig. S4).

### Drug treatments with LDN 193189

LDN-193189 ("LDN") was dissolved to 10mM in dimethyl sulfoxide (DMSO) and stored at -20° C. Fish were live stained with Alizarin Red as described above in groups of ∼24, then split in half and placed into two 2L containers of tank water with light aeration at 18° C for small molecule treatment. Fish were treated with either 20uM LDN in 1% DMSO or 1% DMSO only for three days in parallel. Fish were fed bloodworms during the treatment. After three days, fish were washed with tank water 2x 15 minutes and returned to their 29L tank of origin for 15 days. Fish were fed normally during the rest of the pulse-chase interval. After 18 total days, fish were live stained in calcein as described above, before being fixed for scoring.

### Preparation and blinding of pulse-chase labeled tooth fields

Following fixation, fish from OE or LDN experiments were rinsed and washed in tap water, then agitated in 1% KOH for 20+ minutes at room temperature. Dental tissues were dissected out in tap water or 1x PBS, then washed at room temperature with 1% KOH overnight or with 5% KOH for 30-60 minutes, rendering the mCherry signal no longer detectable, if present. Dental tissues were then washed through a glycerol series (25, 50, 90% glycerol in 1x PBS). Stickleback pharyngeal tissues were flat mounted as previously described (Ellis and Miller, 2016), while zebrafish pharyngeal tooth fields and stickleback oral jaws were arrayed into 24 well plates. The resulting pule-chase dental samples were then shuffled and renamed irrespective of treatment condition so that a different researcher could blindly score and count the teeth in each dental preparation.

### Scoring and analyzing pulse-chase assays

Dental preparations were scored on a Leica M165 stereomicroscope using GFP2, eGFP, and Rhodamine filters to observe Alizarin Red and calcein staining. Every tooth in each animal was addressed for both species (left and right premaxillary, dentary, DTP1, DTP2, and VTP tooth fields in sticklebacks, and the VTP in zebrafish). Importantly, we find that the pulse-chase signal is markedly more difficult to interpret after ∼1 week at room temperature or ∼3 weeks stored at 4° C, both because the calcein signal fades and because autofluorescence in soft tissues increases. Skeletal preparations from a given experiment were thus always scored within two days of the end of each experiment, and always within a single 24 hour window. Since actively growing bony tooth tissues (dentine and enameloid) strongly incorporate these stains, this pulse-chase strategy allows us to classify each tooth in each individual as either “new” or “retained” with respect to the treatment interval: “new” teeth are those that began bone deposition after the Alizarin Red pulse (during the OE treatment), and are thus only marked by the 2^nd^ stain, calcein, while “retained” teeth are those that show any Alizarin Red stain, because this indicates that these teeth were depositing bone prior to the treatment interval (see Fig. 2C and D). The only exception to these rules was in the stickleback *Bmp6* OE treatment, where tooth germs in the treatment condition were observed without either stain. Because these tooth germs were Alizarin Red negative, we inferred that these were new teeth despite their lack of calcein stain. We thus interpret that this class of tooth germs had halted bone deposition by the time the calcein chase occurred. The new:retained ratio and total tooth number for each fish was additionally calculated by dividing or summing new and retained counts, respectively. We additionally counted morphological tooth families in each zebrafish specimen to address primary tooth number expansion or contraction in this species. Since tooth number is known to be dependent on fish size, and oral tooth number is known to be additionally dependent on fish sex (Caldecutt et al., 2001), we used a generalized linear model approach to determine whether corrections for fish sex and/or size were appropriate prior to comparing tooth counts and ratios from transgene carrying and mock heat shock fish groups. The following corrections were applied as per the best fit model: both size (standard length) and sex corrections were applied to *Wnt10a* OE retained and total tooth numbers, *Grem2a* total tooth number, and the *Bmp6* negative control total tooth numbers. Size only corrections were applied to *Grem2a* OE new tooth numbers and area measurements for all four stickleback OE experiments. Sex only corrections were applied to *Dkk2* and *Grem2a* OE retained tooth numbers, and the *Bmp6* negative control retained tooth numbers. No associations between tooth variables and sex or standard length were detected for any of the three zebrafish experiments. All statistical tests and corrections were performed in R (version 2022.02.040+443) (R Core Team, 2022), using two-sided Wilcoxon Rank-Sum tests (Wilcoxon, 1945) in all cases. *P* values were adjusted using the Benjamini-Hochberg method (Benjamini and Hochberg, 1995) to correct the false discovery rate for each experiment where multiple statistical tests were performed on the same groups of individuals. After scoring was complete, example pulse-chase samples were imaged on a Leica DM2500 compound microscope, Leica M165 stereomicroscope, or Zeiss LSM 700 confocal microscope.

### *In situ* hybridization

*In situ* hybridization on sections from subadult and adult sticklebacks was performed as previously described (Square et al., 2021). See Table S2 for probe template sequence information. Probes designed to *Wnt10a* and *Bmp6* were published previously (Cleves et al., 2014; Square et al., 2021). A 3’ UTR probe template for *Grem2a* was cloned from genomic DNA via PCR using the primers GGTGCAGAGGGTCAAACAGT and ATACAGGCTCGTGTCCAAGC. The probe template for *Dkk2* was created from the purchased full-length coding sequence (Gene Universal, Delaware, USA) that was also used to create the overexpression construct (described above). Digoxygenin-labeled *in situ* riboprobes were synthesized as previously described (Square et al., 2021). WT material was embedded in paraffin and sectioned as previously described on either the sagittal or transverse plane (Square et al., 2021).

### Hematoxylin and Eosin staining

*Dkk2* OE and sibling WT fish underwent a typical OE treatment but were not pulsed or chased with Alizarin Red or calcein. These fish were then fixed and sorted as described above. Thereafter these fish were decapitated, and their heads were prepared for sectioning on the transverse plane a previously described (Square et al., 2021). Sections were stained with Hematoxylin and Eosin as previously described (Square et al., 2021).

### RNA-seq

Bulk RNA-seq was performed essentially as described previously (Hart et al., 2018; Mack et al., 2023). 12 hours after a single heat shock, three OE and three control sibling sticklebacks were euthanized in 250 mg/L MS-222 and their VTPs were collected and placed into 50 µl of Trizol (Invitrogen). Samples were shaken by hand, briefly spun down, and incubated on ice for 30-60 minutes to dissociate ribonucleoprotein complexes. Samples were stored at -80° C for up to 3 months. Samples were later thawed on ice, homogenized with a pestle, and extracted with a 1/5th volume of chloroform (10 µl). The resulting supernatant was DNAse treated per standard methods to degrade genomic DNA. Thereafter, the samples were re-extracted with phenol:chloroform and chloroform in Phase Lock Heavy tubes (5PRIME), precipitated with ethanol, and resuspended in 30 µl of DEPC-treated water. Small subsets of each sample were run on a 1% agarose gel and quantified with a Nanodrop to preliminarily assess RNA quality and concentration prior to being analyzed on a Bioanalyzer. After RNA integrity and concentration was determined to be satisfactory, poly-T bead selection, library synthesis, and 100PE Illumina sequencing on a NovaSeq platform were performed by QB3 Genomics, (UC Berkeley, Berkeley, CA, RRID:SCR_022170). Each sample yielded approximately 20M reads. Reads were aligned to the stickleback reference genome (Nath et al., 2021) (version 5; https://ftp.ncbi.nlm.nih.gov/genomes/all/GCF/016/920/845/GCF_016920845.1_GAculeatus_UGA_version5/) with Star (Dobin et al., 2013) using a modified annotation file that included an overexpression transgene sequence as a separate synthetic chromosome in order to sieve out the OE construct transcripts. We removed both the endogenous gene that we overexpressed and the overexpression construct hits prior to differential expression analysis because we could not distinguish between the endogenously expressed gene and the overexpression transgene for most reads aligning to these gene bodies. DEseq2 (Love et al., 2014) was thereafter used in R to test for differential expression of RNAs between the treatment and control conditions.

### Enhancer scoring and imaging

Dental fields from fish carrying one copy of the TCF/Lef reporter transgene (Square et al., 2021) were prepared and analyzed essentially as described previously (Stepaniak et al., 2021) with some modification. Tooth fields were dissected from euthanized fish and fixed lightly in 4% formaldehyde at room temperature for 5 minutes on heavy agitation, washed through a glycerol series (5 minutes each in 25, 50, 90% glycerol in 1x PBS), mounted between glass coverslips, and scored and/or imaged on a Leica DM2500 compound microscope.

## Supporting information

Table S1: probe template sequences

Table S2: overexpression transgene sequences

---

**Figure S1.**
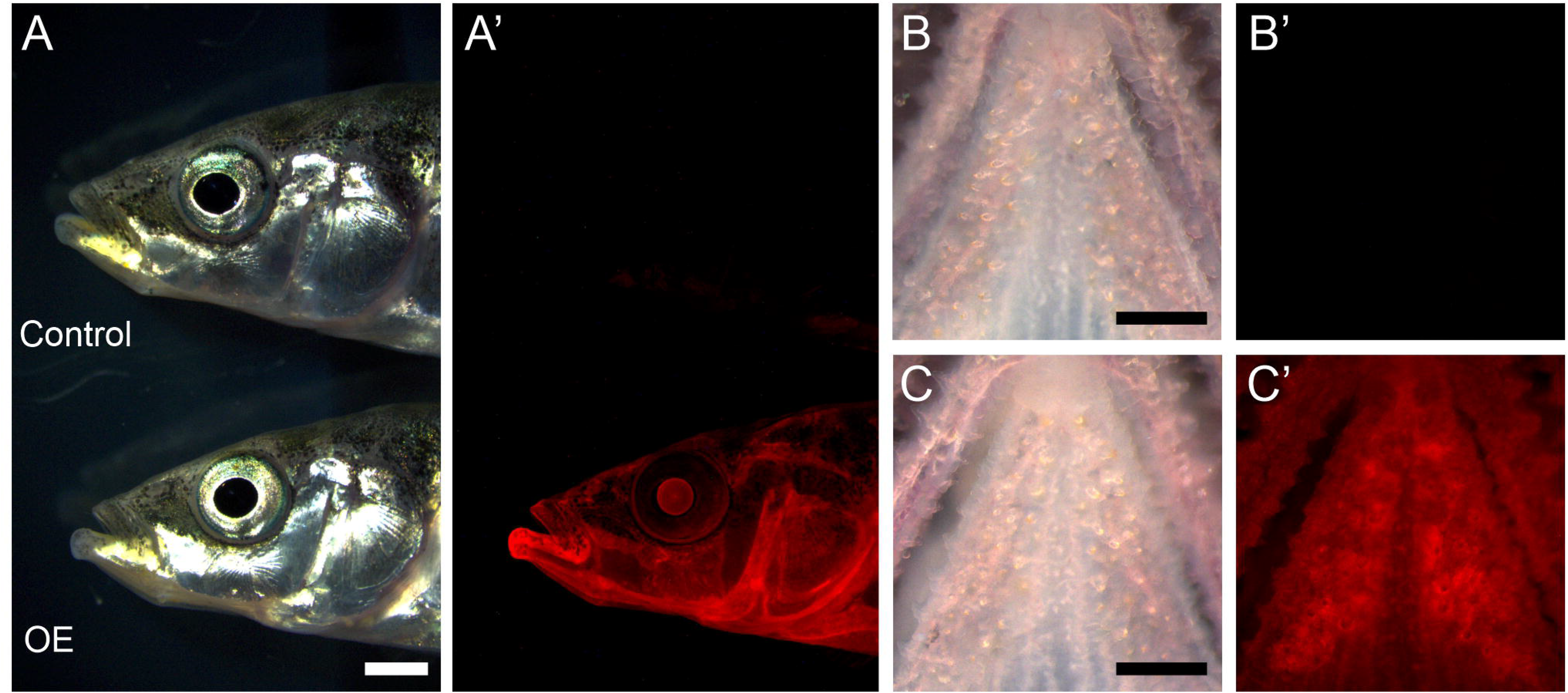
Zebrafish *hsp70l* activation in stickleback. **A.** Lateral images of sticklebacks 24 hours after a single heat shock. Control (sibling without transgene) on top, overexpression transgene positive (OE) fish on bottom (*Bmp6* OE in this example). C’ shows red channel fluorescence, allowing visualization of the transgene. **B and C.** Dissecting the ventral tooth plate (VTP) from control, OE fish revealed mCherry present throughout the tooth field (B’ and C’ show red channel fluorescence), anterior to top. Scale bar in A: 2 mm, B and C: 500 μm.

**Figure S2.**
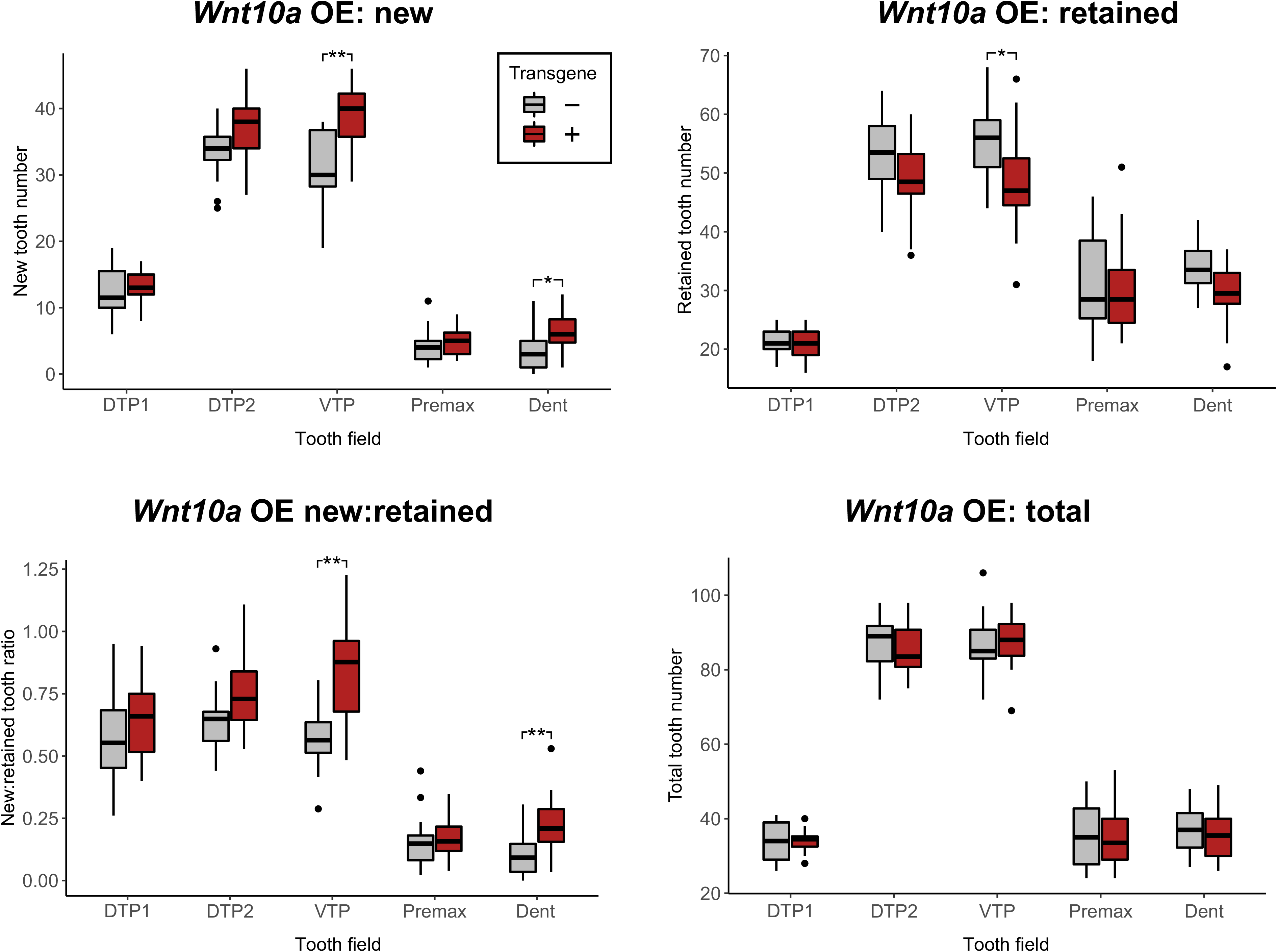
Results of stickleback *Wnt10a* OE parsed by tooth field reveals strongest results on ventral tooth fields (ventral tooth plate and dentary). Stickleback pharyngeal teeth are housed in three discrete fields: the 1^st^ dorsal tooth plate 1 (DTP1), the 2^nd^ dorsal tooth plate (DTP2), and the ventral tooth plate (VTP). In the oral jaws, sticklebacks have teeth on their premaxilla (Premax) and dentary (Dent) bones. Each group of graphs shows new, retained, the new:retained ratio, and total tooth number broken down by tooth field type (the sum of left and right halves per fish). \**P*<0.05, \*\**P*<0.01, \*\*\**P*<0.001 (Wilcoxon Rank-Sum tests, Benjamini-Hochberg corrected *P* for multiple hypothesis testing). Insignificant results (*P*>0.05) are unannotated. Boxes represent the 25th-75th percentiles, and the median is shown as a gray bar.

**Figure S3.**
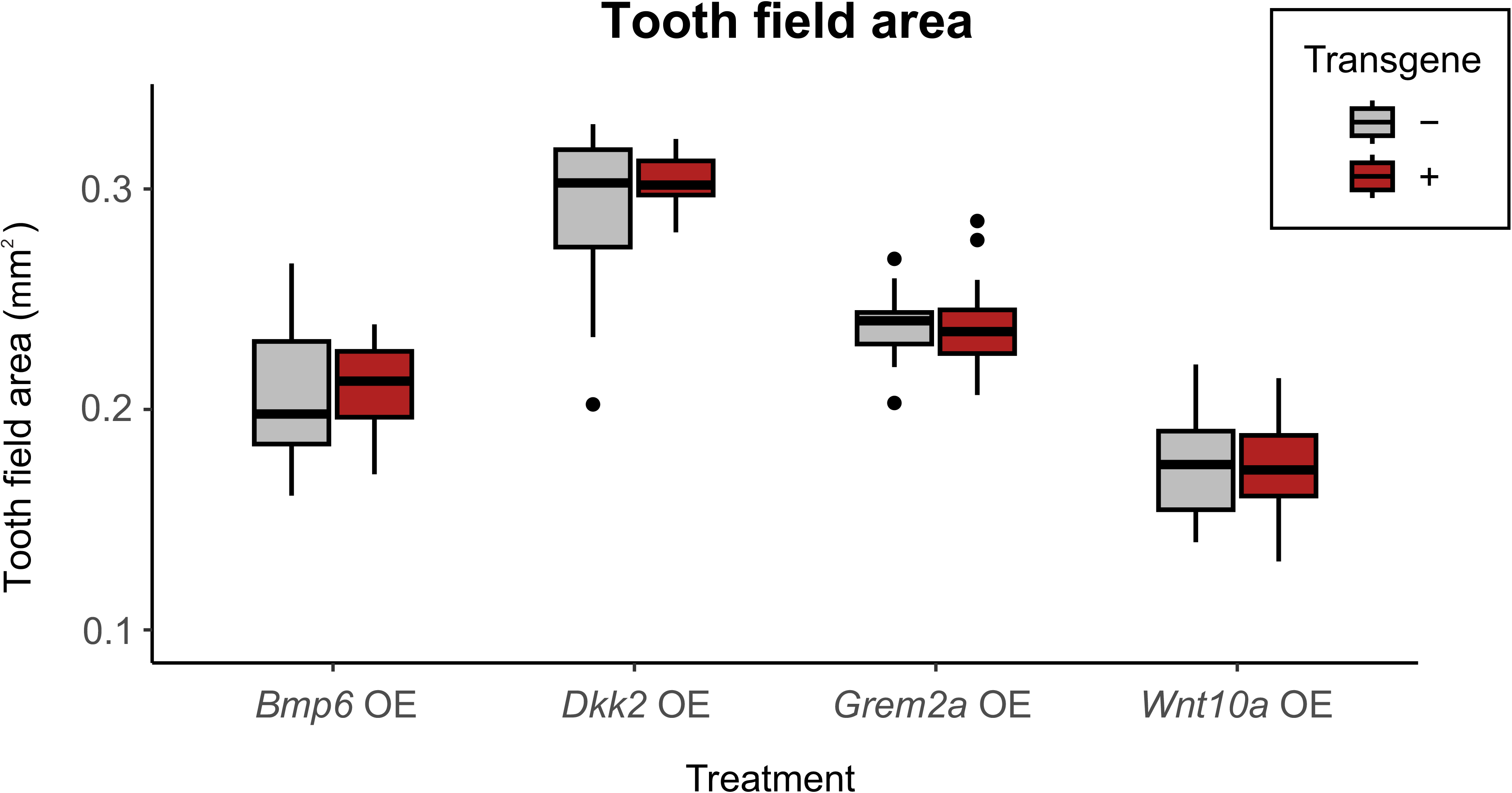
VTP area did not demonstrate significant changes under any OE experiment. Box and whisker plots showing left VTP area (mm^2^) for all sticklebacks that underwent a heat shock treatment. Area values were corrected for standard length. No significant differences were observed in any experiment (all *P*>0.4, Wilcoxon Rank-Sum tests). Boxes represent the 25th-75th percentiles, and the median is shown as a gray bar.

**Figure S4.**
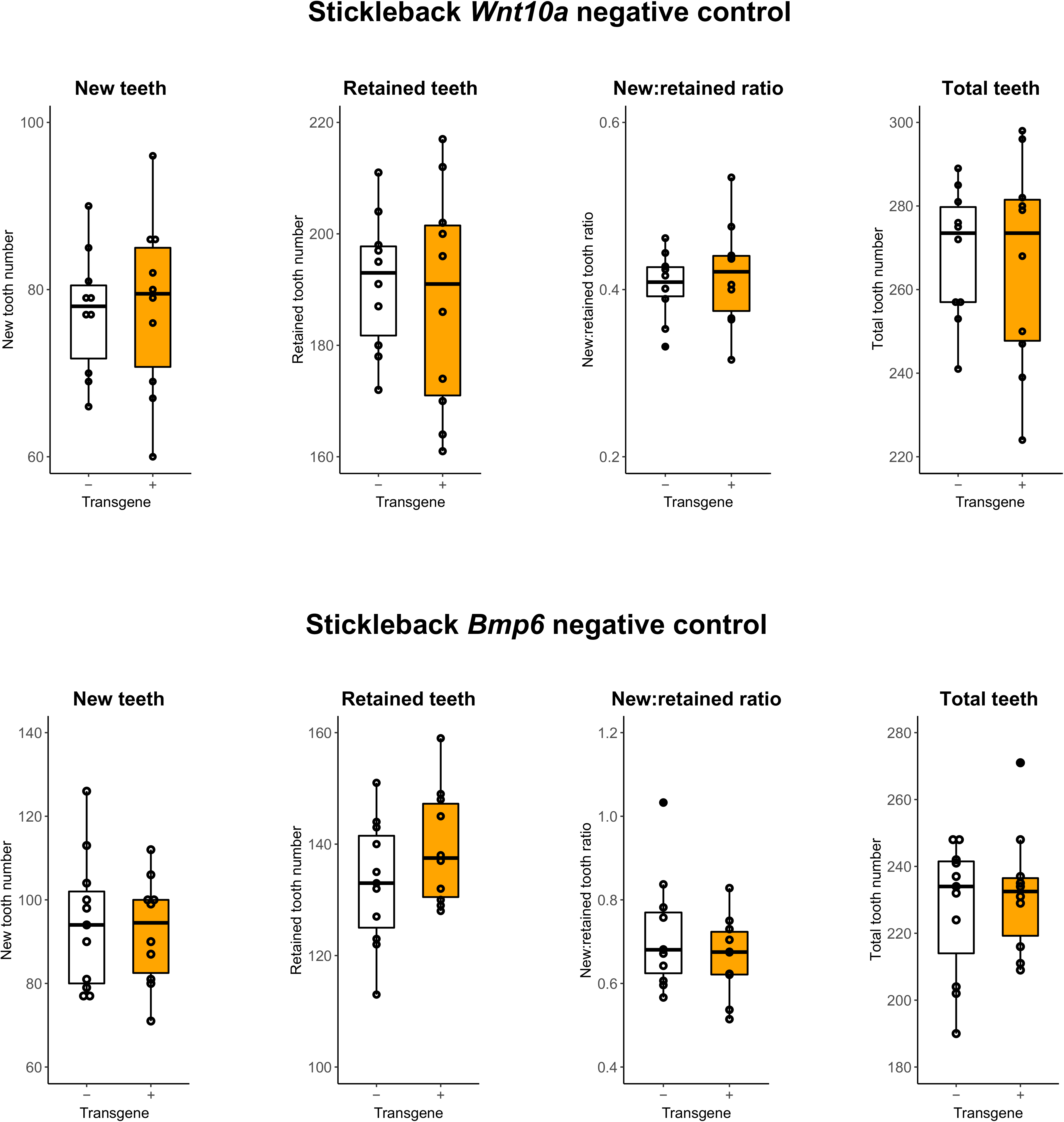
Negative control experiments suggest inducible transgenes in the absence of heat shocks have little to no effect on tooth replacement dynamics. The same pulse-chase interval (18 days, see Methods) with no intervening heat shocks was applied to full siblings of the fish in the experiments summarized in Fig. 3A and Fig. 6A, testing for whether simply carrying the transgene in the absence of any heat shock is associated with changes in the number of new teeth, retained teeth, the new:retained ratio, or total teeth; no such deviations were detected in this negative control experiment for any of these four variables assessed (all *P*>0.90 and *P*>0.26 for *Wnt10a* and *Bmp6* negative control experiments, respectively, Wilcoxon Rank-Sum tests, Benjamini-Hochberg adjusted for multiple hypothesis testing. No “stalled” tooth germs were observed in either negative control assay. Boxes represent the 25th-75th percentiles, and the median is shown as a gray bar.

**Figure S5.**
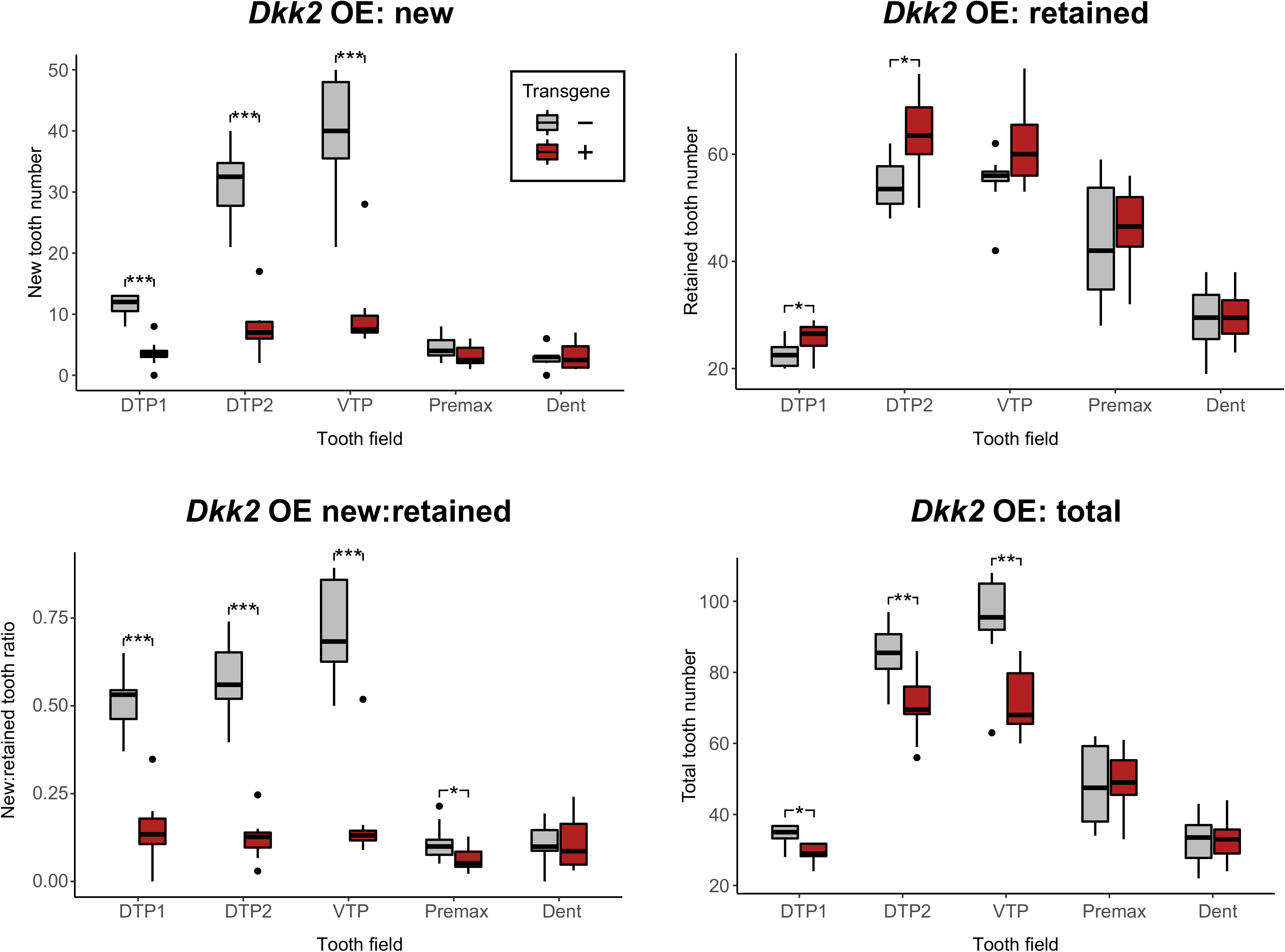
Results of stickleback *Dkk2* OE parsed by tooth field reveals effects primarily on pharyngeal teeth. Stickleback pharyngeal teeth are housed in three discrete fields: the 1^st^ dorsal tooth plate 1 (DTP1), the 2^nd^ dorsal tooth plate (DTP2), and the ventral tooth plate (VTP). In the oral jaws, sticklebacks have teeth on their premaxilla (Premax) and dentary (Dent) bones. Each group of graphs shows new, retained, the new:retained ratio, and total tooth number broken down by tooth field type (the sum of left and right halves per fish). \**P*<0.05, \*\**P*<0.01, \*\*\**P*<0.001 (Wilcoxon Rank-Sum tests, Benjamini-Hochberg corrected *P* for multiple hypothesis testing). Insignificant results (*P*>0.05) are unannotated. Boxes represent the 25th-75th percentiles, and the median is shown as a gray bar.

**Figure S6.**
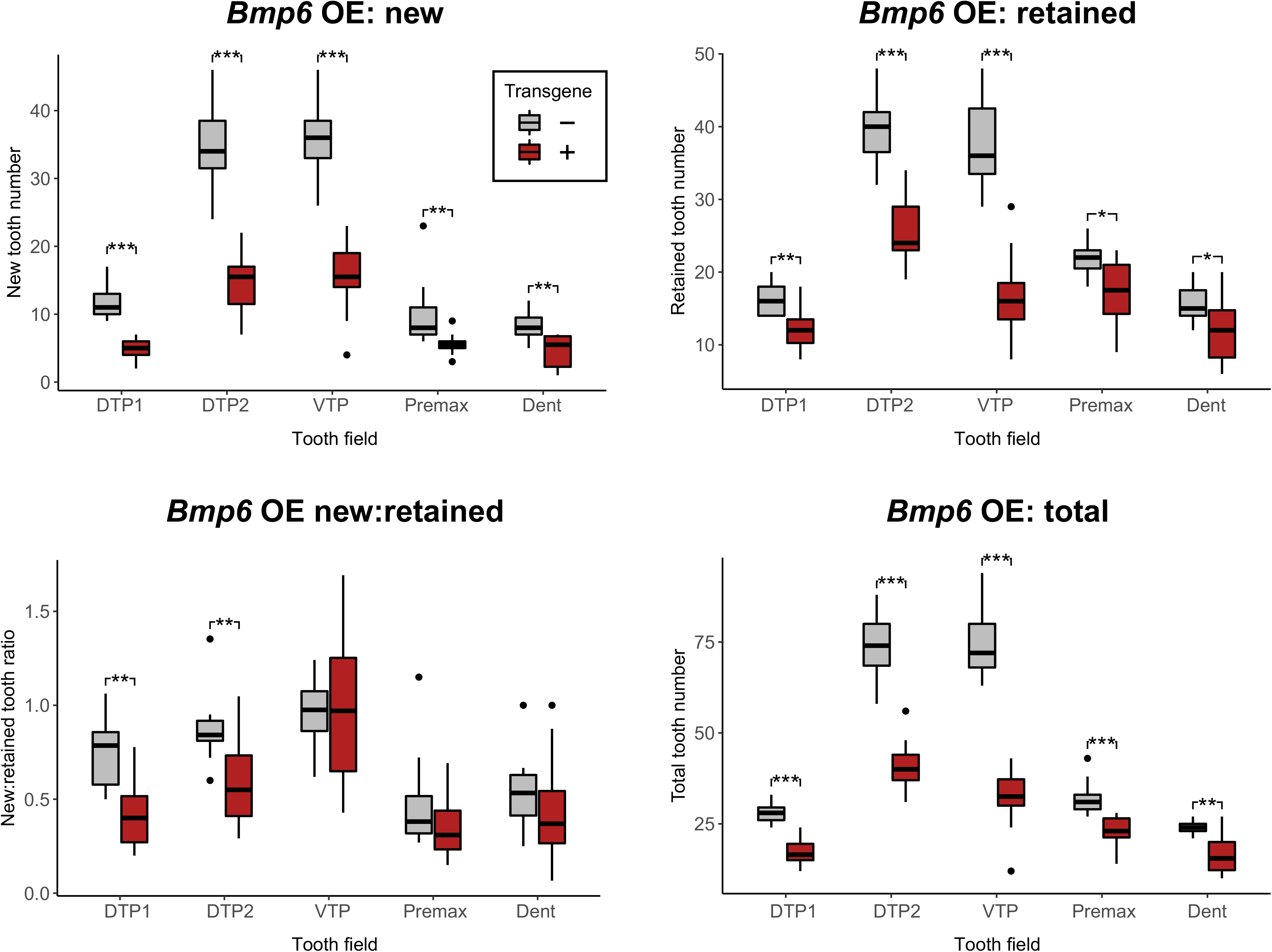
Results of stickleback *Bmp6* OE parsed by tooth field reveals more pronounced effects on pharyngeal teeth than oral teeth. Stickleback pharyngeal teeth are housed in three discrete fields: the 1^st^ dorsal tooth plate 1 (DTP1), the 2^nd^ dorsal tooth plate (DTP2), and the ventral tooth plate (VTP). In the oral jaws, sticklebacks have teeth on their premaxilla (Premax) and dentary (Dent) bones. Each group of graphs shows new, retained, the new:retained ratio, and total tooth number broken down by tooth field type (the sum of left and right halves per fish). \**P*<0.05, \*\**P*<0.01, \*\*\**P*<0.001 (Wilcoxon Rank-Sum tests, Benjamini-Hochberg corrected *P* for multiple hypothesis testing). Insignificant results (*P*> 0.05) are unannotated. Boxes represent the 25th-75th percentiles, and the median is shown as a gray bar.

**Figure S7.**
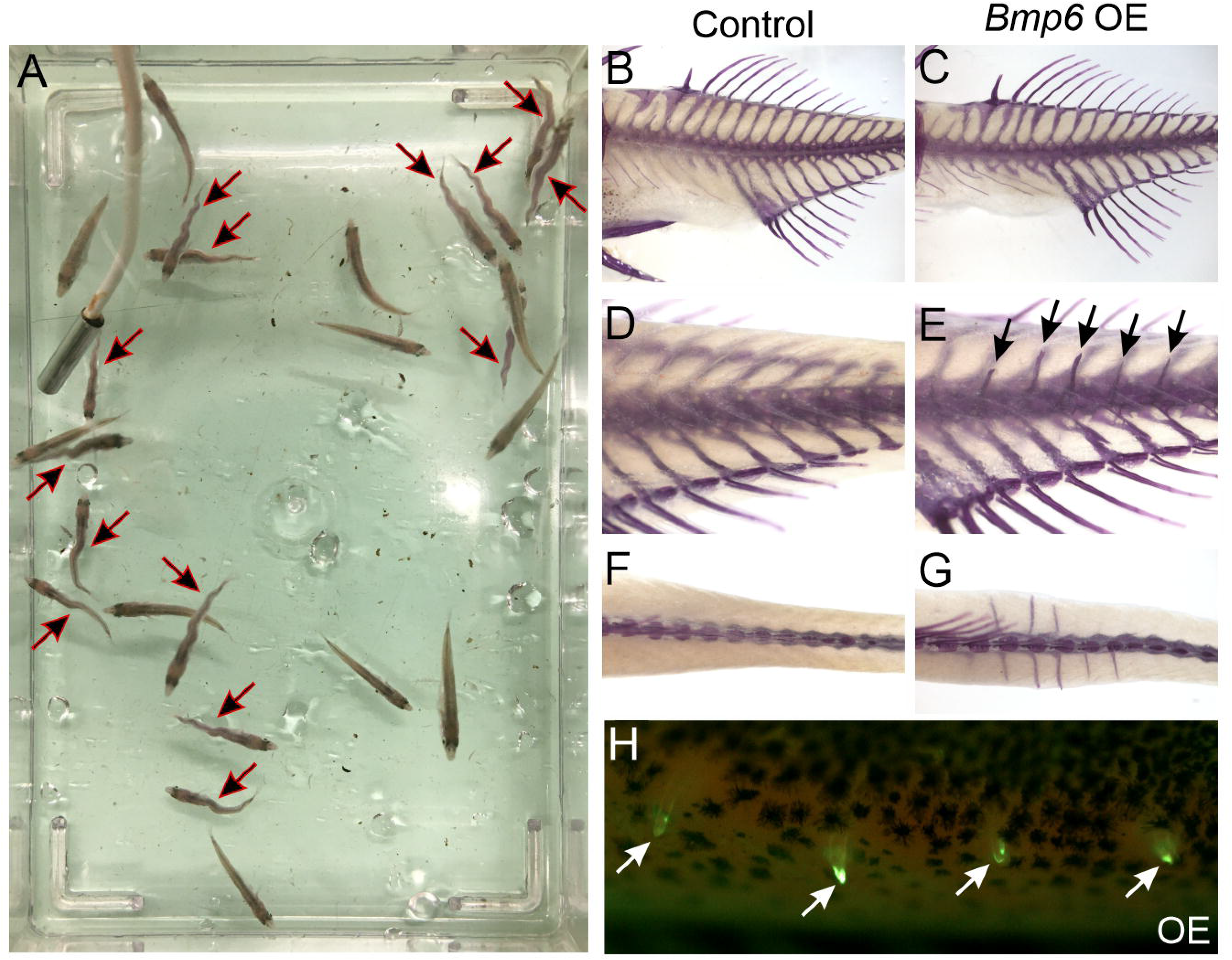
Axial bending and ectopic rib-like bony protrusions from *Bmp6* OE. 14/14 OE fish had bends along their primary axis (arrows in A), 8/14 had some number of bony body spikes (black arrows in C, white arrows in D), all of which were confined to caudal vertebrae 3-8. B-G show fish skeletons that were re-stained with Alizarin Red after the pulse-chase experiment to beter visualize the skeleton. B and C show straight lateral view, D and E show oblique lateral views tilted ∼30 degrees on the coronal axis, F and G show dorsal views. H shows a lateral image of a treatment animal immediately after the pulse-chase experiment, before re-staining, revealing that the new bone growth is indeed strongly marked by calcein, confirming that these body spikes arose during the OE treatment interval.

**Figure S8.**
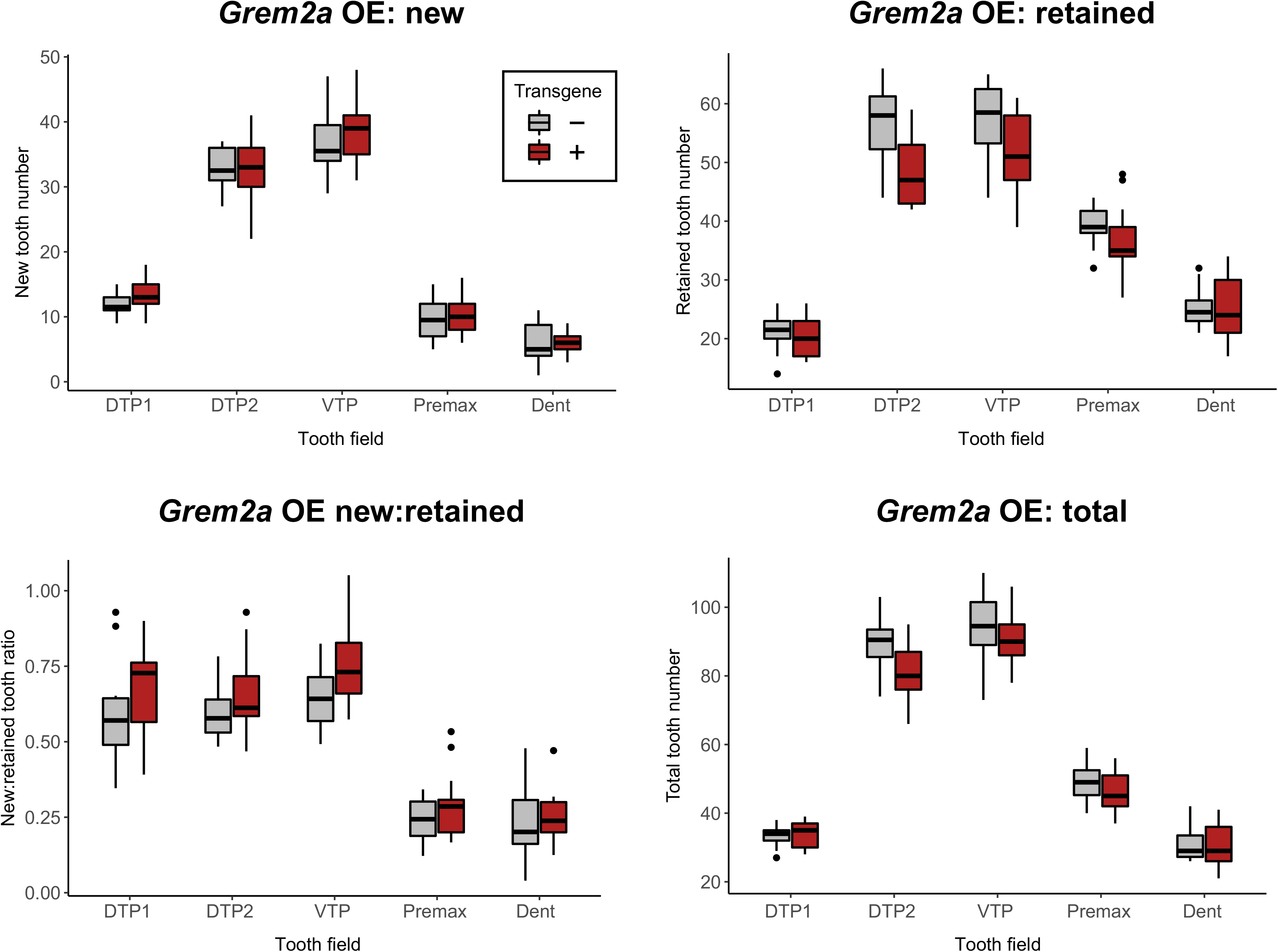
Results of stickleback *Grem2a* OE parsed by tooth field reveals consistent but subtle effects on new, retained, and the new:retained ratio. Stickleback pharyngeal teeth are housed in three discrete fields: the 1^st^ dorsal tooth plate 1 (DTP1), the 2^nd^ dorsal tooth plate (DTP2), and the ventral tooth plate (VTP). In the oral jaws, sticklebacks have teeth on their premaxilla (Premax) and dentary (Dent) bones. Each group of graphs shows new, retained, the new:retained ratio, and total tooth number broken down by tooth field type (the sum of left and right halves per fish). All *P*>0.1279 (Wilcoxon Rank-Sum tests, Benjamini-Hochberg corrected *P* for multiple hypothesis testing). Boxes represent the 25th-75th percentiles, and the median is shown as a gray bar.

**Figure S9.**
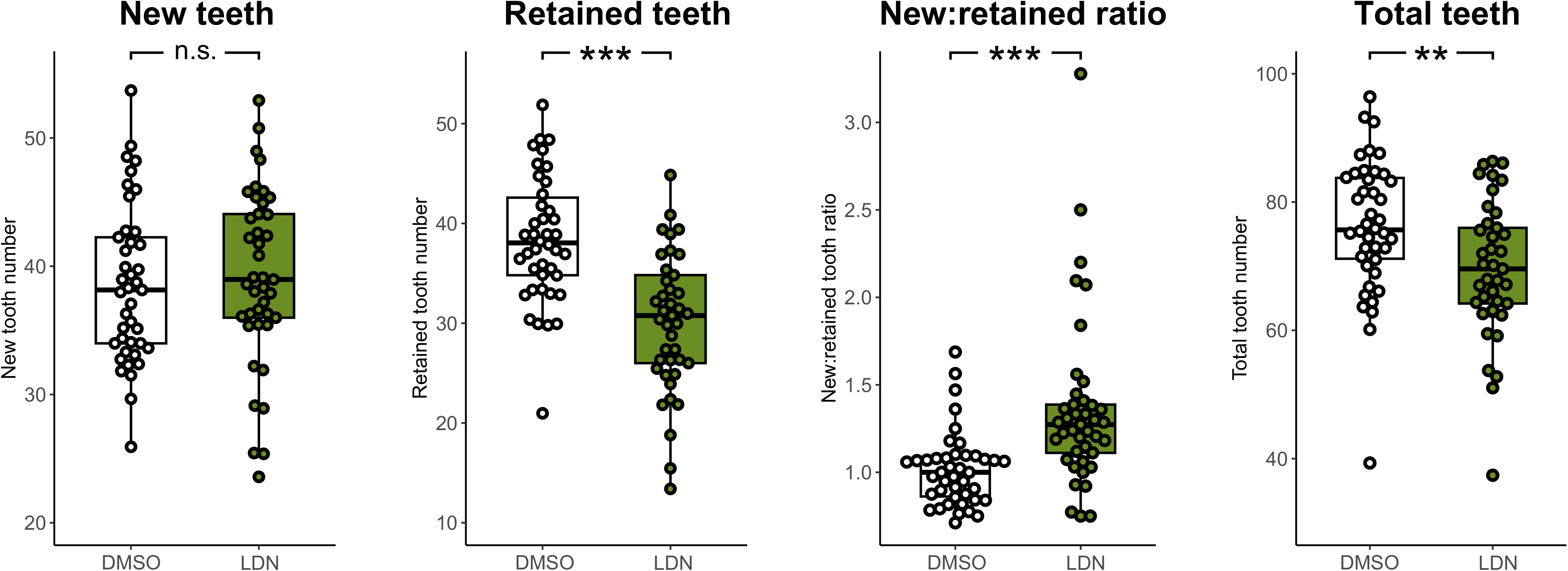
Effects of the BMP signaling inhibitor LDN 193189 on ventral tooth plates (VTP). Using the same pulse-chase bone-labeling approach as in the OE experiments, we inhibited BMP signaling via LDN 193189 (n=42 control and 41 treatment fish). New tooth number did not significantly change (Wilcoxon Rank-Sum Benjamini-Hochberg adjusted *P*=0.34), retained teeth significantly decreased (*P*=4.9e-6), the new:retained tooth ratio significantly increased (*P*=1.5e-5), and total number of teeth decreased (*P=*0.0055). Boxes represent the 25th-75th percentiles, and the median is shown as a gray bar.

## Acknowledgements

We thank Andrew Glazer for cloning the stickleback *Grem2a* riboprobe template, Tess Linden and Nicole King for sharing the pMS48 plasmid, Mark Stepaniak for creating and sharing the T2Rv10 plasmid, Shane Schwikert for *pro bono* statistics consultation, Diana Aguilar Gomez and Katya Mack for assistance with RNA-seq software and analysis, and Sophie Archambeault, Dennis Sun, and Claire Altier for feedback on the manuscript. This work was supported by NIH grants DE027871, DE031017, and DE021475.

## Notes

### Competing Interest Statement

The authors have declared no competing interest.

### Summary of Updates

We perfomed additional experiments to assay the genetic response to Bmp6 overexpression, finding an upregulation of Wnt inhibitors, a downregulation of Wnt10a, and reduced expression of a TCF/Lef reporter under. We othrwise improved the statistical analyses, and streamlined the text.

